# An Adipo-Pulmonary Axis Mediated by FABP4 Hormone Defines a Therapeutic Target Against Obesity-Induced Airway Disease

**DOI:** 10.1101/2024.07.15.603433

**Authors:** M. Furkan Burak, Gurol Tuncman, Ayse Nur Ayci, Kashish Chetal, Grace Yankun Lee Seropian, Karen Inouye, Zon Weng Lai, Nurdan Dagtekin, Ruslan I. Sadreyev, Elliot Israel, Gökhan S Hotamışlıgil

## Abstract

Obesity-related airway disease is a clinical condition without a clear description and effective treatment. Here, we define this pathology and its unique properties, which differ from classic asthma phenotypes, and identify a novel adipo-pulmonary axis mediated by FABP4 hormone as a critical mediator of obesity-induced airway disease. Through detailed analysis of murine models and human samples, we elucidate the dysregulated lipid metabolism and immunometabolic responses within obese lungs, particularly highlighting the stress response activation and downregulation of surfactant-related genes, notably SftpC. We demonstrate that FABP4 deficiency mitigates these alterations, demonstrating a key role in obesity-induced airway disease pathogenesis. Importantly, we identify adipose tissue as the source of FABP4 hormone in the bronchoalveolar space and describe strong regulation in the context of human obesity, particularly among women. Finally, our exploration of antibody-mediated targeting of circulating FABP4 unveils a novel therapeutic avenue, addressing a pressing unmet need in managing obesity-related airway disease. These findings not only define the presence of a critical adipo-pulmonary endocrine link but also present FABP4 as a therapeutic target for managing this unique airway disease that we refer to as fatty lung disease associated with obesity.

**Graphical Abstract:** 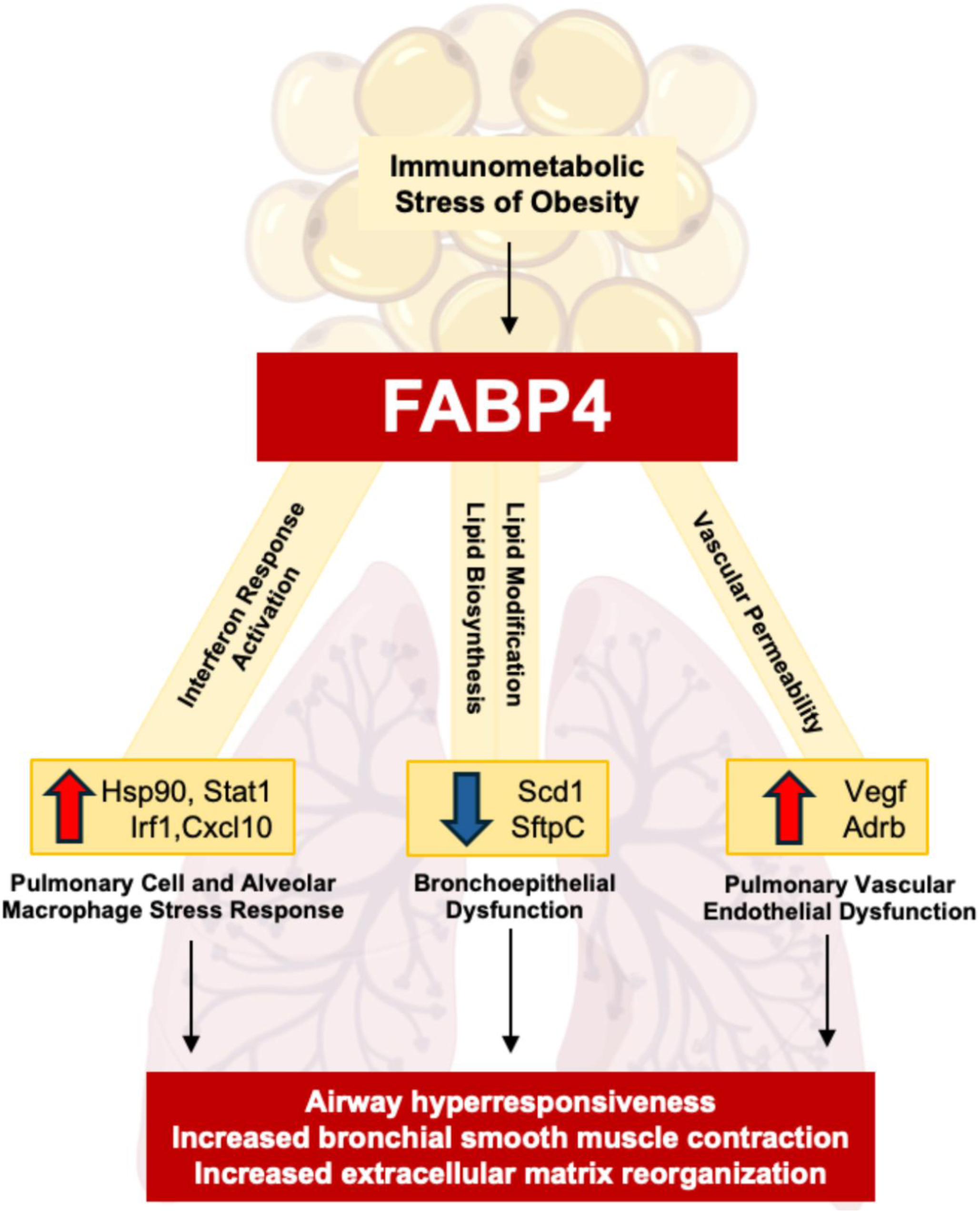

**One Sentence Summary:** Investigating FABP4’s pivotal role in obesity-driven airway disease, this study unveils an adipo-pulmonary axis with potential therapeutic implications.

## Introduction

Obesity has a profound negative impact on the pulmonary system, including heightened susceptibility to infections and complications, due to immunometabolic dysregulation, which was highlighted during the SARS CoV-2 pandemic. However, the ramifications of obesity on respiratory function extend far beyond infection susceptibility. Thoracic fat deposition, for instance, constricts lung expansion, while increased truncal obesity flattens the diaphragm. Consequently, compensatory respiratory efforts often prove inadequate, leading to poor central respiration signals resulting in weakened accessory respiratory muscles and obesity-driven hypoventilation in some cases. This imbalance contributes to increased blood flow to the poorly ventilated lung base, creating a ventilation-perfusion mismatch, thereby altering airway dynamics, reducing inspiratory capacity, and inducing rapid, shallow breathing. However, it is also clear that the adverse effects of increased adiposity on pulmonary function are far beyond its mere restrictive mechanical impact. These alterations may potentially foster airway hyperresponsiveness and dysfunction (*1, 2*). As adipose tissue expands, it recruits various immune cells. Along with adipocytes, these cells collectively secrete adipokines, chemokines, and cytokines. This metabolic inflammation, termed “metaflammation,” lies at the core of many obesity-related pathologies, encompassing diabetes, fatty liver disease, and intriguingly, a subset of asthma (*3*). Consequently, obesity-related airway disease is often used synonymously with asthma, although a clear definition and characterization of this condition remain incomplete.

Asthma has evolved into an umbrella term encompassing reactive airway diseases. While classic T-helper type 2 (Th2)-driven allergic/eosinophilic asthma, influenced by genetic predisposition and environmental triggers, is well-documented, several other asthma phenotypes also exist. Asthma with obesity, on the other hand, presents a much more complex and unusual pattern. Albeit oversimplified, epidemiological studies delineate two categories of airway disease linked to obesity: 1) Early-onset allergic asthma exacerbated by obesity and 2) Late-onset non-allergic asthma developing as a complication of obesity (*4–7*). The latter primarily affects women, exhibits low inflammatory markers, does not feature eosinophilia, and often shows improvement in airway symptoms with weight loss. Individuals with obesity experience a more rapid progression in the severity of asthma, with frequent exacerbations necessitating emergency care, and inadequate responses to conventional treatments, often succumbing to their side effects such as steroid related hyperglycemia, osteoporosis, and adrenal insufficiency (*8, 9*). Unfortunately, novel therapeutic approaches, such as biologics mostly address patients with Type 2 inflammation (*10*), while effective treatments are not available for patients with obesity-related airway disease.

Immunometabolic processes contribute to asthma endotypes. Lipid mediators, which play a significant role in airway inflammation and hyperresponsiveness, show upregulation in obesity (11, 12). This positions obesity not only as a major risk factor for airway disease but also as a potential disease modifier (*6, 11–13*). Notably, approximately 60% of severe asthma patients exhibit obesity (*14–16*), suggesting a potential intersection in the chronic immuno-metabolic pathways of these two conditions. The lipid chaperone fatty acid binding protein 4 (FABP4) is an exceptionally versatile immunometabolic regulator in both intra and intercellular context (*17, 18*). This 15 kDa protein possesses a lipid-binding pocket that orchestrates intracellular lipid trafficking and signaling, while also regulating metabolic and inflammatory responses. For example, deletion of FABP4 in macrophages blocks inflammatory cytokine production and related gene expression (*19, 20*), whereas in adipocytes, FABP4 is coupled to lipid breakdown and stress signals (*17*). These properties are critical in FABP4’s role in the pathogenesis of metabolically driven chronic low-grade inflammatory diseases such as obesity, diabetes, fatty liver disease, and atherosclerosis, as firmly established in the literature (*17, 21, 22*).

Recent findings revealed the secretion of FABP4 from adipocytes and endothelial cells into circulation, presenting it as an endocrine hormone significantly elevated in obese mice and humans (*23, 24*). Elevated FABP4 levels correlate strongly with worsened metabolic, inflammatory, and cardiovascular outcomes across multiple independent human studies (*25–34*). Notably, individuals carrying a polymorphic allele causing reduced FABP4 expression exhibit a decreased risk of developing diabetes, dyslipidemia, and cardiovascular disease, particularly in overweight and obese subjects (*28, 31, 35*). Targeting circulating FABP4 with a monoclonal antibody (mAb) has demonstrated efficacy in alleviating metabolic dysfunctions in preclinical models of diabetes, fatty liver disease, and obesity (*36, 37*). These cumulative findings underscore FABP4’s conservation of biological functions relevant to human pathophysiology and presenting circulating FABP4 as a validated and promising therapeutic target in humans (*17, 21, 22*). Interestingly, FABP4 has also been linked to allergic asthma and respiratory infections (*38, 39*), although it is not known whether this activity relates to local or systemic exposure to the hormonal form of this protein.

In this study, we characterized the obese lung pathology in detail and documented the unique features of this condition. Our systematic unbiased analyses and transcriptional profiles revealed regulatory patterns consistent with FABP4 function. Our single-cell RNA sequencing also unveiled pathways through which obesity primes the lungs toward various diseases, including but not limited to asthma. Interestingly, in addition to its increased levels in the circulation, we elucidated the presence of FABP4 in the bronchoalveolar space in an obesity-regulated manner, both in obese mice and humans, and linked its elevation to airway hyperresponsiveness. Through transplantation studies, we established adipose tissue as the source of FABP4 in the alveolar space, uncovering a unique adipo-pulmonary endocrine link. The genetic deletion of the FABP4 protected mice from airway hyperresponsiveness in both genetic and diet-induced obesity models. Crucially, targeting circulating FABP4 via an anti-FABP4 monoclonal antibody significantly ameliorated airway hyperresponsiveness in severely obese mice. These data support FABP4 as a pivotal mediator in this distinctive form of obesity-associated airway disease, where increased FABP4 levels from adipose tissue integrate metabolic status with lung biology. These findings highlight FABP4 not only as a marker for a subset of human asthma associated with obesity but also as a promising target for developing new therapeutics to address a significant medical gap.

## Results

### Obesity causes intrapulmonary fat accumulation and primes the lungs for airway disease

Given the profound impact of obesity on susceptibility to various lung diseases, exploring the underlying alterations in obese lungs and airways becomes crucial. Our investigation started with a detailed macroscopic and microscopic examination of the chest cavity in obesity. Transverse plane visualization exhibited substantial fat deposition in the mediastinum of genetically obese mice (Fig. 1A), prominently surrounding the heart and ultimately extending into the chest cavity. Computed tomography confirmed increased mediastinal and subcutaneous fat in obese mice, potentially contributing to restricted inspiratory capacity without any visible parenchymal damage (Fig. 1B). Intriguingly, microscopic analysis revealed an increased intrapulmonary fat, which was confirmed with perilipin staining (right bottom image in Fig. 1C). This fat was notably concentrated around the larger airways and vessels, accounting for approximately 2.5% of lung sections-, around five times more than in lean lung tissue (Fig. 1C, Fig. S1A and B). This observation aligns with previous findings in postmortem lungs of obese humans, with and without asthma, further substantiating our results (*40*).

**Figure 1.**
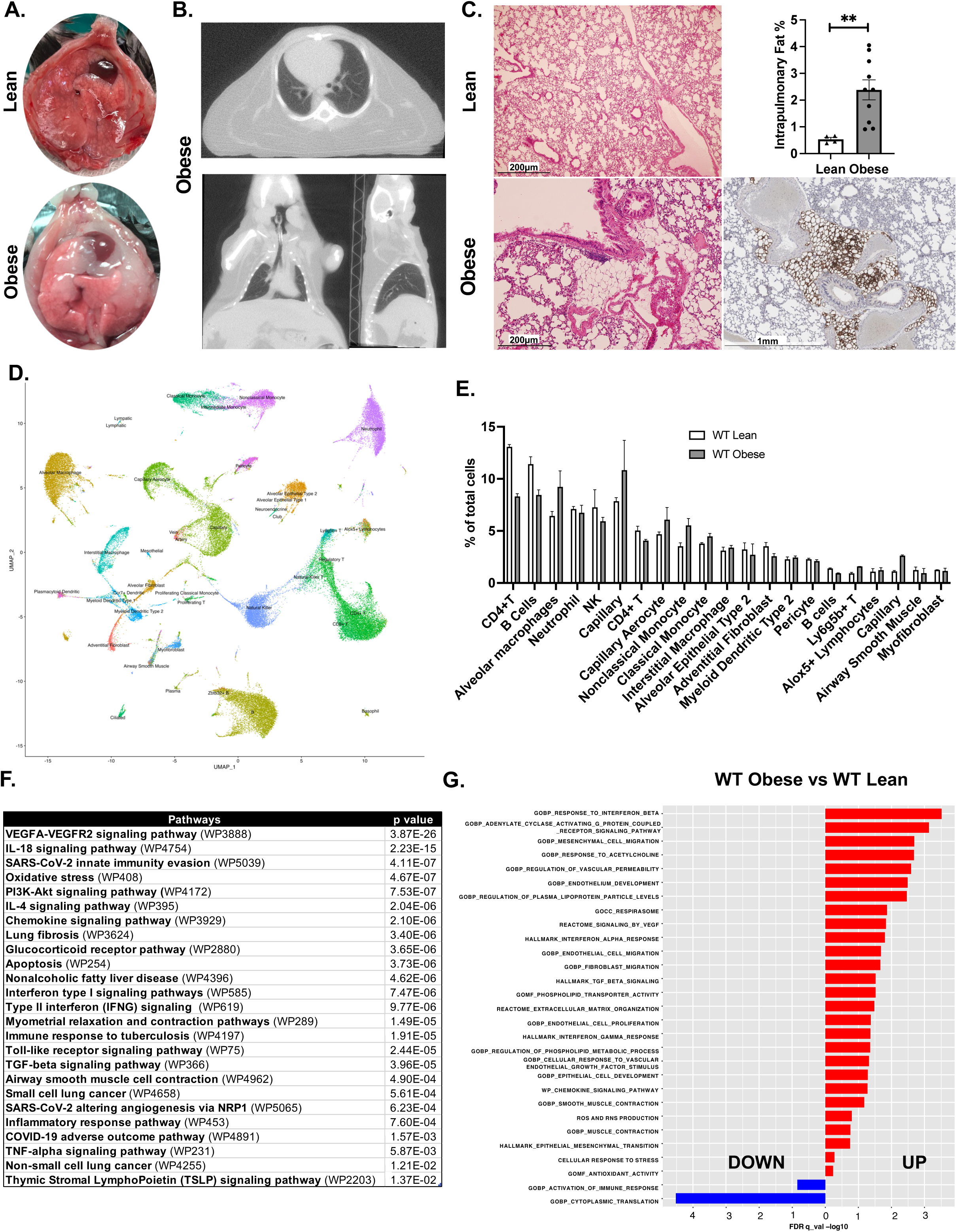
Macroscopic and molecular characterization of obese lungs. **A.** Macroscopic images of the thoracic cavities of lean and obese (ob/ob) mice, transverse section. **B.** CT images of the thoracic region of an obese mouse. Clockwise from the top: Images from transverse plane, sagittal plane, and coronal plane. **C.** Microscopy images of the lungs from lean (top left) and obese (bottom left) mice with H&E staining, fat cells stained with perilipin (bottom right), and quantification of the intrapulmonary fat area (top right). Fat area calculated as a percentage of the adipocyte area to the whole lung section area. n=4-10/group. **D.** Single-cell RNA-seq analysis of cell preparations from whole lungs of lean and obese mice (18000-20000 cells/group). We used a UMAP (Uniform Manifold Approximation and Projection) plot, analyzed by the ELeFHAnt tool, to annotate the predicted total of 41 cell type clusters using mouse cell atlas data as a reference (79). **E.** Cell clusters from the lungs of lean and obese mice, the most abundant 21 cell clusters are plotted. **F.** Pathway enrichment analysis of the 1,385 differentially expressed genes from the 21 most abundant cell clusters in lean and obese mice, using the Enrichr tool (WikiPathway 2021 Human). The top pathways altered in obese lungs, ranked by p-value. **G.** Top enriched relevant pathways in WT obese vs. WT lean mice, identified using GSEA on whole lung tissue (all 41 cell clusters).

To comprehensively understand molecular alterations associated with obesity, we conducted single-cell RNA sequencing and identified 41 distinct clusters in the obese lung (Fig. 1D). Contrary to our expectation that a single cell type would explain the obese phenotype, we observed alterations across multiple cell types, such as reductions in T-, B-, and NK cells, epithelial cells, fibroblasts, and airway smooth muscle cells, alongside increases in capillary cells, aerocytes, monocytes, and alveolar macrophages (Fig. 1E). EnrichR analysis of functional category enrichment (*41*) among the differentially expressed genes (DEGs) in the twenty most abundant clusters identified several pathways altered in obese lungs, such as VEGFA-VEGFR2, IL-4, Toll-like Receptor, and TNF-α signaling. Interestingly, most of these top enriched WikiPathway (*42*) categories were also linked to FABP4 action in obesity (*17, 20, 38, 43–46*) (Fig. 1F). Gene set enrichment analysis (GSEA) (*47*) across all 41 cell clusters in the whole lung highlighted the upregulation of pathways associated with *bona fide* asthma features, such as increased acetylcholine response, smooth muscle contraction, extracellular matrix organization, and remodeling pathways in obese lungs. (Fig. 1G). However, obesity-related asthma is characterized by the absence of the allergic T helper 2 (Th2) phenotype, meaning there is no eosinophilic airway inflammation. Instead, it is accompanied by inflamed adipose tissue, metabolic imbalance, and dyslipidemia—features also observed in obese mice (*3*). Additionally, fat deposition in and around the lungs and upregulated pathways indicate a unique form of airway phenotype, that we propose to refer as the fatty lung disease associated with obesity.

### FABP4 deficiency protects lungs from obesity-induced stress response

Since GSEA analysis revealed dramatic upregulation in both known FABP4 downstream pathways (*20, 36, 38, 43–46, 48*) and asthma gene signatures (Fig. 1G), we examined the potential impact of FABP4 in these obesity-related alterations directly. We made a comparative evaluation of these targeted pathways between FABP4-deficient (FABP4 KO) obese vs. wildtype (WT) obese lungs. Strikingly, this analysis revealed that a significant proportion of the obesity-induced changes in expression patterns across various functional gene categories were reversed in the lung tissue of FABP4 KO mice (Fig. 2A and B).

**Figure 2.**
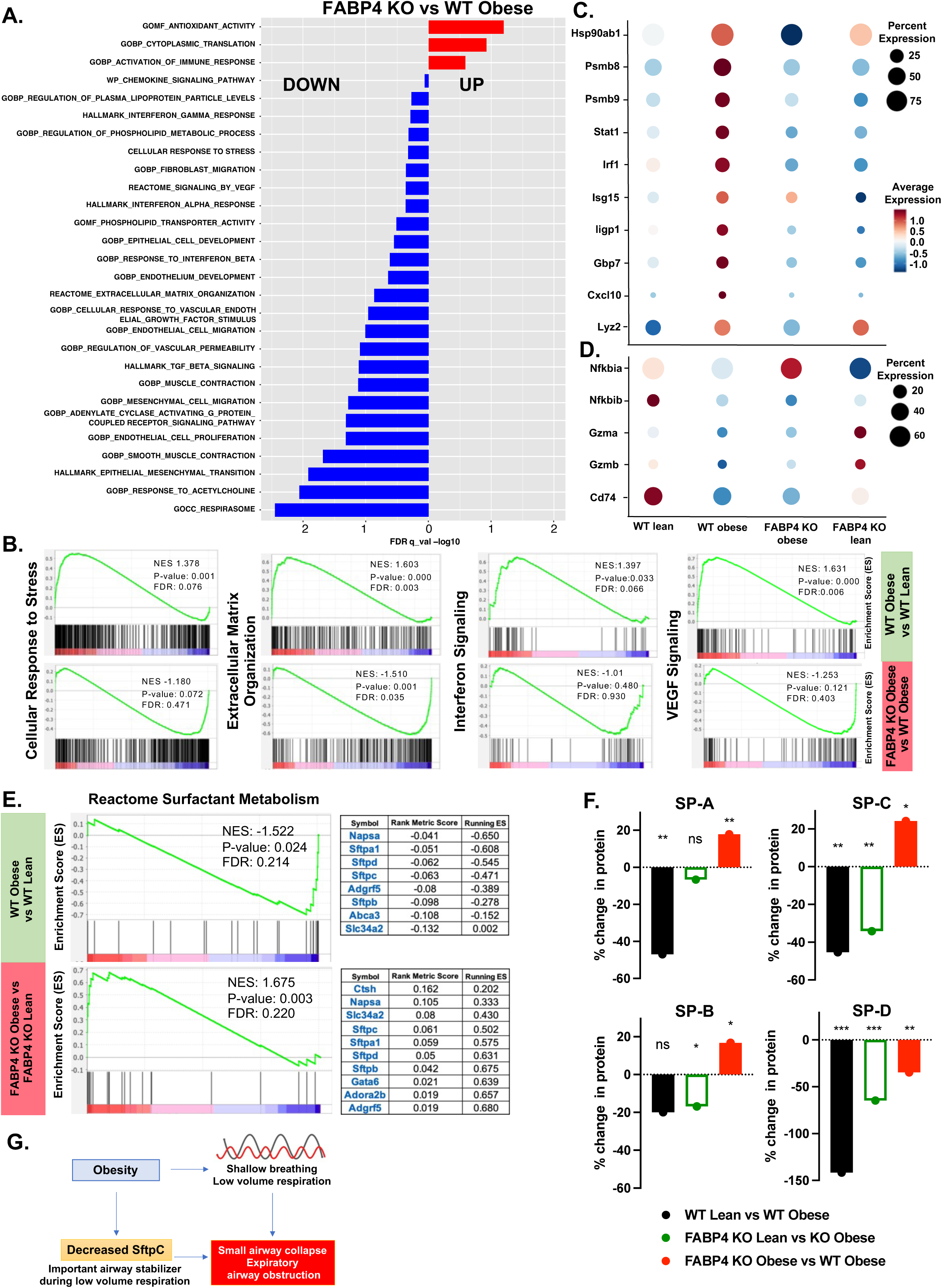
Activation of stress response pathways and decreased surfactant in obese lungs. **A.** Top enriched relevant pathways (same pathways from figure 1G) in FABP4-deficient (FABP4 KO) obese vs. wild type (WT) obese mice, identified using GSEA on whole lung tissue (all 41 cell clusters). **B.** Representative enrichment plots from GSEA analysis. **C.** Dot plot expression analysis of the genes involved in stress response and interferon response pathways, and **D.** Genes protecting from stress response across all clusters from WT lean, WT obese, FABP4 KO obese and FABP4 KO lean lungs, respectively. **E.** GSEA plots of surfactant metabolism pathway genes, with core enriched genes in the type 2 epithelial cell clusters comparing WT obese vs. WT lean and FABP4 KO obese vs. FABP4 KO lean. On the right side of the GSEA plots, only core enriched genes are listed by rank order. **F.** Measurements of surfactant proteins (SP-A, SP-B, SP-C, and SP-D) from BALF, performed by proteomics, shown as percent differences in absolute protein amounts between WT lean vs. WT obese, FABP4 KO lean vs. FABP4 KO obese, and FABP4 KO obese vs. WT obese. **G.** Working model of obesity-related airway obstruction through decreased surfactant protein C (SftpC).

Focusing on stress response pathways, we observed a dramatic upregulation of Hsp90 expression in the lungs of obese WT mice compared to lean ones. Notably, this upregulation was absent in the lungs of obese FABP4 KO mice, indicating that the absence of FABP4 prevents the activation of these pathways (Fig. 2C). Further exploration revealed the broad activation of the interferon response pathway (Stat1, Irf1, Isg15, Gbp7, Iigp1, Cxcl10) in WT obese lungs (Fig. 2C), which was accompanied by downregulation of protective genes (Nfkbia, Nfkbib, Gzma, Gzmb, Cd74) against stress response (Fig. 2D). These alterations are strikingly mitigated in FABP4 KO obese lungs, indicating FABP4’s pivotal role in regulating pulmonary stress response pathways in obesity (Fig. 2C and D). *In vitro* validation studies in an alveolar macrophage cell line treated with recombinant FABP4 showed increased release of Vegf, Gdf-15, M-csf, interferon-gamma and downstream mediators such as Cxcl10 and Cxcl11, affirming the influence of FABP4 in stress response pathways (Fig. S2A).

The stress response also affected surfactant and related genes, which is another key component of the functional and healthy alveolar space. Surfactant, secreted by alveolar epithelial cells, not only protects against airway collapse by reducing alveolar surface tension but also contributes to various immunometabolic functions (51). Multiple studies have shown that surfactant proteins are downregulated in asthma, and recombinant surfactant treatment improves airway hyperresponsiveness (*49–52*). Interestingly, consistent with the literature, GSEA in the alveolar epithelial cell cluster revealed that the surfactant pathway was downregulated in WT obese lungs compared to lean, but not in FABP4 KO mice, paralleling the expression of surfactant genes (Fig. 2E). To measure the amount of surfactant proteins in the alveolar space, we performed proteomics analysis of the bronchoalveolar lavage fluid (BALF) samples collected from the same groups, which further confirmed reduced surfactant proteins in WT obese compared to lean controls. Strikingly, FABP4 KO obese mice had significantly higher levels of surfactant proteins (SP-A, SP-B, and SP-C) (Fig. 2F) than WT obese mice. Shallow breathing and low-volume respiration in obesity can cause the collapse of small airways, potentially leading to airway obstruction, which has been proposed as one of the mechanisms of obesity-related asthma (1). Given the fact that surfactant proteins are key stabilizers of airways during low-volume respiration (56, 57) (Fig. 2G), the downregulation of SftpC in obesity can exacerbate airway collapse and hyperresponsiveness.

Since surfactant is a complex molecule composed of approximately 10% protein and 90% lipid, we also analyzed the lipid-related pathways. GSEA in epithelial cells revealed downregulation of genes involved in lipid biosynthesis and modification in obesity (Fig. S2B). Specifically, genes such as Scd-1, Fas, ApoE, Acly, Fads1, Elovl1, and Cd36 showed altered expression in obese lung epithelial cells, indicating disrupted lipid metabolism (Fig. S2B). In our cohort, the Scd-1 gene, a key player in lipogenesis, was exclusively expressed in epithelial cells, but its expression was significantly decreased in obesity. On the other hand, the fatty acid transporter Cd36 gene expression was upregulated in all clusters of cells in obese lungs (Fig. S2C), suggesting differential effects of increased lipid exposure in different cell clusters. Our findings align with previous studies showing that decreased Scd1 expression is associated with reduced surfactant protein C levels in bronchoepithelial cells of humans with asthma compared to healthy controls, and it contributes to airway hyperresponsiveness in mice (58).

Additionally, we identified upregulated inflammatory pathways in alveolar macrophages, increased vascular permeability-related pathways in endothelial cells, and decreased surfactant proteins, alongside altered lipid biosynthesis pathways in epithelial cells. These findings suggest that multiple cell clusters, particularly those surrounding the alveolar space, contribute to obesity-related changes underlying obese asthma phenotype (Fig. 1G and Fig. S1C). Notably, FABP4 deficiency protects against these alterations in obese lungs (Fig. 2A). However, it remains unclear whether local or systemic FABP4 mediates this marked effect under these conditions.

### FABP4 levels increase systemically and locally in the airways of obese mice and humans with obesity

The multifaceted alterations observed through single-cell RNA sequencing and proteomics in obese lungs were notably absent in genetic FABP4 deficiency. Examination of Fabp4 expression revealed minimal signal in type 2 bronchoepithelial cells, capillary cells, monocytes, and alveolar macrophages (Fig. 3A), with no notable changes in expression patterns or intensity between obese and lean conditions (data not shown). These observations prompted us to explore the potential endocrine impact of FABP4 on lung physiology. Initially, we assessed FABP4 levels in plasma from lean and obese mice, confirming elevated levels in both genetic and diet-induced obesity models compared to lean controls (Fig. 3B, S3A). Subsequently, we collected BALF to investigate local airway presence of FABP4. Surprisingly, we not only detected FABP4 in the bronchoalveolar space but also observed a significant upregulation in obese mice (Fig. 3C, S3B).

**Figure 3.**
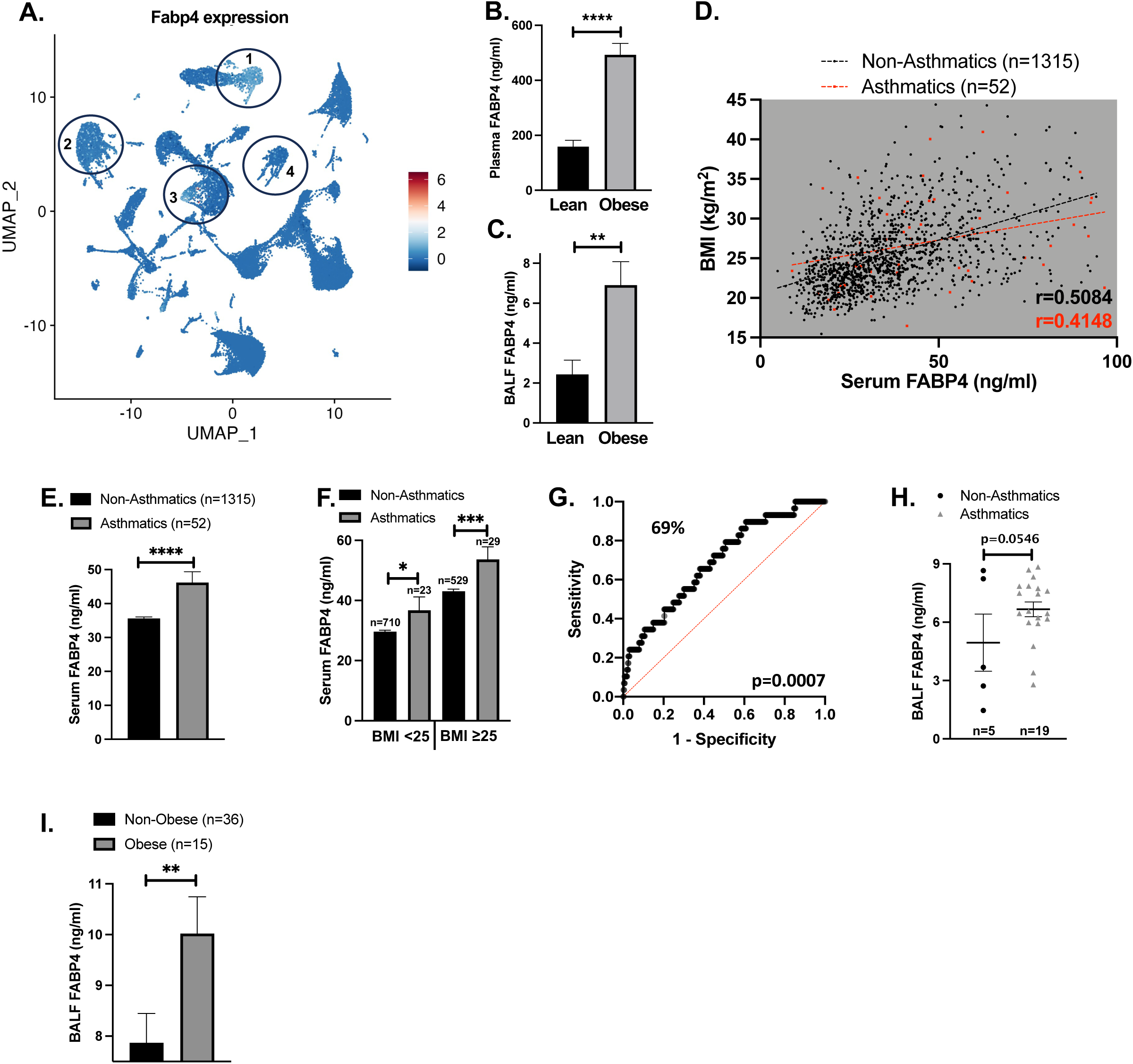
FABP4 levels in serum and BALF are increased in obese mice and humans. **A.** FABP4 gene expressing clusters marked on UMAP plot of the whole lung.1- Non-classical monocytes, 2- Alveolar macrophages, 3- Endothelial cells, 4- Broncho-epithelial cells. **B.** Plasma FABP4 levels and **C.** BALF FABP4 levels, measured by ELISA, n=16-23/group, from lean (WT) and obese mice (ob/ob, genetic obesity). **D.** Serum FABP4 levels from female humans with and without asthma. Correlation of BMI and serum FABP4 levels calculated using the Spearman test. r = 0.5084, p < 0.0001 in the non-asthmatic group, and r = 0.4148, p = 0.0021 in the asthmatic group. **E.** Serum FABP4 levels from females with and without asthma, **F.** Serum FABP4 levels in females with and without asthma, stratified by lean (BMI < 25) vs. overweight/obese (BMI ≥ 25) status. Sample sizes were 710 and 23 for lean females without asthma and with asthma, respectively. In the overweight/obese groups, sample sizes were 592 and 29 for females without asthma and with asthma, respectively. **G.** Receiver operating characteristic curve (ROC) predicting asthma status based on serum FABP4 levels. p= 0.0007. **H.** BALF FABP4 levels from women with and without asthma (p=0.0546), each symbol represents one person. **I.** BALF FABP4 levels from obese vs. non-obese humans, p<0.01 (both males and females).

To determine if our findings in obese mice are applicable to humans, we investigated serum FABP4 levels in individuals randomly selected from the Nurses’ Health Study and Health Professionals Follow-up Study, both with and without asthma. Obesity-related asthma, a unique subtype within the asthma spectrum, is characterized by distinct features. Unlike other subtypes, this form of asthma typically develops later in life and is more prevalent in women. Our data, consistent with earlier reports, showed a positive correlation between serum FABP4 levels and BMI (Fig. 3D). When stratified by asthma status, we observed significantly higher serum FABP4 levels (30% elevation) in females who reported having a physician-diagnosed asthma (*53*) (Fig. 3E). Notably, when stratified by BMI, asthma prevalence was approximately 3.14% (23 out of 733) among lean females (BMI < 25) and about 5.20% (29 out of 621) among overweight and obese females (BMI ≥ 25), representing over a 60% increase in prevalence in our cohort. Additionally, the association between FABP4 levels and asthma status was more pronounced, with a lower p-value, among overweight and obese females compared to lean females (BMI < 25), p<0.05 vs. p<0.001 (Fig. 3F). These findings underscore FABP4 as a potential independent risk factor for obesity-related asthma. Further investigations in large asthma cohorts are needed to validate these findings and distinguish obesity-related subtype. Nonetheless, even in this relatively small study encompassing heterogeneous asthma types, a notable ROC Curve of 0.69 (Fig. 3G) suggests the potential utility of serum FABP4 levels as a biomarker for obesity-related asthma in females.

In our animal studies, we detected FABP4 in BALF, consistent with elevated levels observed in plasma from obese mice (Fig. 3 B and 3C). To determine the relevance of these findings in human disease, we also measured FABP4 levels in BALF collected during elective bronchoscopy from individuals with and without obesity. Obesity-related asthma features a low Th2 involvement and a non-eosinophilic pathology, predominantly affecting females (*11, 14, 54*). Despite sample size limitations, our observations suggest a trend towards FABP4 regulation in BALF samples from females with asthma (p=0.0546) (Fig. 3H). In contrast, we did not observe significant differences in FABP4 levels in males, either in serum (Fig. S3C-E) or BALF (Fig. S3F). Nevertheless, irrespective of asthma status, FABP4 was consistently present in human BALF, with levels 30% higher (p=0.01) in individuals with obesity compared to non-obese controls (Fig. 3I). These findings indicate an increase in FABP4 levels locally within the airways in obese individuals and highlights its potential as a pathological mediator and suggests a direct link between adipose tissue and the lungs, particularly in the context of obesity.

### Adipose tissue-derived FABP4 reaches the airways and creates an adipo-pulmonary axis

Elevated FABP4 levels in the alveolar space in obesity prompted us to inquire about the origin of FABP4 protein. Our prior research demonstrated a substantial increase in FABP4 secretion from adipose tissue during obesity (*24*), leading us to explore whether FABP4 derived from adipose tissue could reach the airways. To examine this, we first conducted an adipose tissue transplantation experiment using WT obese mice as donors and FABP4 KO lean mice as recipients. Adipose tissue from obese WT mice was transplanted into the intraperitoneal region of FABP4 KO mice (Fig. 4A). Following surgery, the mice remained healthy and maintained their body weight (Fig. S4A). Three days post-surgery, we collected blood and BALF samples. Interestingly, ELISA results revealed detectable levels of FABP4 in both the plasma (Fig. 4B) and the BALF in all recipient FABP4 KO mice (Fig. 4C).

**Figure 4.**
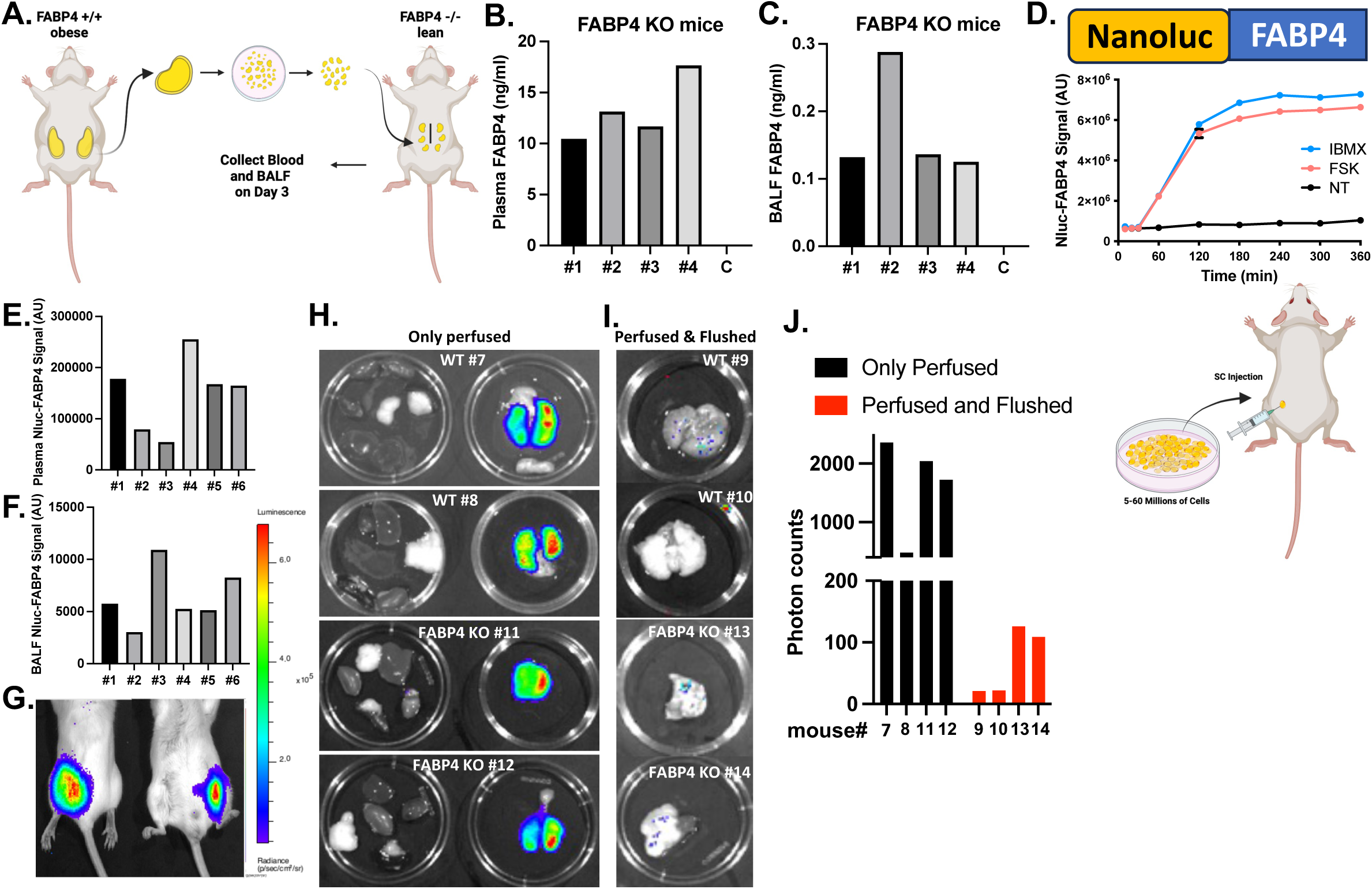
Adipo-pulmonary axis via FABP4. **A.** Adipose tissue transplantation 3-day protocol. **B.** Plasma FABP4 levels and **C.** BALF FABP4 levels from individual recipient FABP4 KO mice (mice #1-4) and a non-transplanted control mouse (C) on day 3 after transplantation, measured by ELISA. **D.** Validation of FABP4 secretion during lipolysis in NanoLuc (Nluc)-FABP4- expressing 3T3-L1 mouse adipocytes, and protocol for adipocyte implantation in mice. NanoLuc signal was quantified using a luciferase assay on conditioned media collected post-treatment with IBMX (1-methyl-3-isobutylxanthine), FSK (Forskolin), or no treatment (NT – vehicle). **E.** Luciferase assay measurements of Nluc-FABP4 signal in plasma and **F.** BALF, collected three days after adipocyte implantation into recipient WT mice (#1-6). **G.** Whole mouse IVIS bioluminescence imaging. Nluc signal measured 30 minutes after substrate injection, exclusively detected at the cell implantation site. **H.** Nluc-tagged FABP4 signal detected in organs (liver, heart, kidney, inguinal fat, spleen, pancreas—left panel) and whole lung (right panel) from recipient WT and FABP4 KO mice, three days after implantation with Nluc-tagged-FABP4 expressing 3T3-L1 adipocytes. Organs were harvested 30 minutes post-substrate injection following perfusion with 20 ml saline. **I.** Image of lungs flushed by lavage with a total of 5 ml saline after whole mouse perfusion. **J.** Quantification of luminescent signal from lungs, plotted as total luminescence count per lung in radiant photons.

To trace the secretion of FABP4 from adipocytes and enhance detection sensitivity, we utilized the 3T3-L1 mouse pre-adipocyte cell line transfected with luciferase-tagged (N-terminal) FABP4. After confirming Nluc-FABP4 secretion via *in vitro* lipolysis (Fig. 4D), we implanted Nluc-FABP4-expressing differentiated adipocytes into both wild type and FABP4 KO mice. Three days post-implantation we collected blood and bronchoalveolar lavage fluid. Using a luciferase assay, we detected luciferase-tagged FABP4 in the plasma in a dose-dependent manner (Fig. S4B). Strikingly, we also observed FABP4 in the BALF (Fig. S4C). The Nluc-signal intensity was higher in WT mice compared to FABP4 KO mice, consistent with our previous observations that FABP4 clearance rates are higher in FABP4 KO mice (unpublished). In a separate FABP4 KO-only cohort, we performed *in vivo* lipolysis, confirming that implanted adipocytes remained healthy and secreted FABP4 into both the circulation and the bronchoalveolar space (Fig. 4E and F). A longer-term analysis showed that the Nluc signal returned to baseline levels 180 minutes after lipolysis induction, confirming the viability of adipocytes after implantation (Fig. S4D). Lastly, simultaneous measurement of FABP4 by luciferase assay and FABP4 ELISA in the same plasma collected from FABP4 KO mice ruled out possible artefact signaling from luciferase system (Fig. S4E and F).

After validating the proper responses with the NanoLuc construct and establishing the number of cells to be transfected, we performed additional experiments in another cohort of WT and FABP4 KO mice. We had three groups: mice injected the NanoLuc-FABP4-expressing cells, mice injected with NanoLuc only expressing cells, and sham mice injected only saline to the dorsal hindleg as a control. Three days after implantation, we injected furimazine, an imidazopyrazinone substrate for NanoLuc (Nluc), into the tail vein of mice and imaged thirty minutes later with *in vivo* optical tomography IVIS® Spectrum (PerkinElmer). We detected the Nluc signal exclusively at the injection site (dorsal and ventral planes) (Fig. 4G). We confirmed that there was no signal in sham control mice (without cells) after substrate injection, nor in transplanted mice (with cells) before substrate injection (Fig. S4G). Since we demonstrated that the implanted adipocytes release Nluc-FABP4 into circulation, we tracked the FABP4 flux to specific organs. To do this, we perfused the heart from the left ventricle with 20 ml saline 30 minutes after the tail vein injection of the substrate. After perfusing the mice and harvesting organs, the substrate alone did not produce any signal (Fig. S4H), and only negligible background signal was detected in organs harvested from Nluc-only adipocyte-implanted mice (Fig. S4I, lower panel). Importantly, we detected a significant Nluc-FABP4 signal only in the lungs. There was no detectable signal in other organs, including but not limited to the kidneys, pancreas, spleen, epididymal fat, and brain (Fig. 4H). This suggests the presence of FABP4 in the non-vascular compartments of the lungs, either in the parenchyma or bronchoalveolar space. Due to the air-blood (alveolar-capillary) barrier, perfusing the lung through the heart does not affect the contents of the alveolar space. To further determine the Nluc-FABP4 protein location, we lavaged the airways of perfused mice with 5 ml saline by flushing through the trachea to clear the bronchoalveolar space. Strikingly, we observed an almost complete disappearance of the Nluc-FABP4 signal, confirming that the FABP4 signal in the lungs of perfused mice was emerging from the bronchoalveolar space (Fig. 4I). In addition to analyzing signal distribution, we also quantified the emitted photons. The perfused and flushed lungs showed a significantly lower signal compared to lungs that were only perfused, where alveolar content remained intact (average photon counts: 69 vs. 1653, respectively), further supporting our conclusions. (Fig. 4J). Collectively, these experiments strongly validate the existence of a previously unrecognized adipo-pulmonary link mediated by adipose tissue-derived FABP4.

### FABP4 deficiency or targeting circulating FABP4 via a monoclonal antibody attenuates obesity-related airway hyperresponsiveness independent of adiposity

As obesity exacerbates airway hyperresponsiveness in humans (*1*) and our single-cell RNAseq data revealed upregulated asthma-relevant pathways in the obese lungs, we hypothesized that obesity would induce airway hyperresponsiveness in mice, and lack of FABP4 would alleviate the effects of obesity on the airways. To test our hypothesis, we first assessed the airway reactivity of severely obese (ob/ob) mice to increasing doses of the bronchoconstrictive agent methacholine (MeCh), without any allergic or environmental (i.e., ozone) challenge. Since FABP5 expression is upregulated in adipose tissue in the absence of FABP4 but not regulated in obesity (*24*), we decided to first test our hypothesis in double KO (FABP4 and FABP5 double knockout – FABP deficient) mice to avoid any potential masking or confounding effect of FABP5 upregulation and isolate the role of FABP4. Strikingly, obese (ob/ob) mice exhibited dramatically higher airway resistance without any allergen challenge, whereas littermate FABP- deficient ob/ob mice had markedly reduced airway resistance related to obesity, despite no effect of FABP deletion on body weight (Fig. 5A and B). Encouraged by these results, we further validated our hypothesis using a short-term high-fat diet model in FABP4-deficient (FABP4 KO) mice. We observed increased airway resistance in WT obese mice compared to lean WT, whereas FABP4-deficient obese mice exhibited attenuated resistance, independent of body weight (Fig. S5A and B). These consistent findings across independent obesity models underscore FABP4’s crucial role in obesity-related airway hyperresponsiveness. From the airway physiology experiments described above, we achieved a better window in our bronchoprovocation tests with ob/ob mice. For this reason, we chose to proceed with the genetic model of obesity, which eliminates the potential confounding effects of leptin and age, as these mice gain weight consistently and at earlier stages.

**Figure 5.**
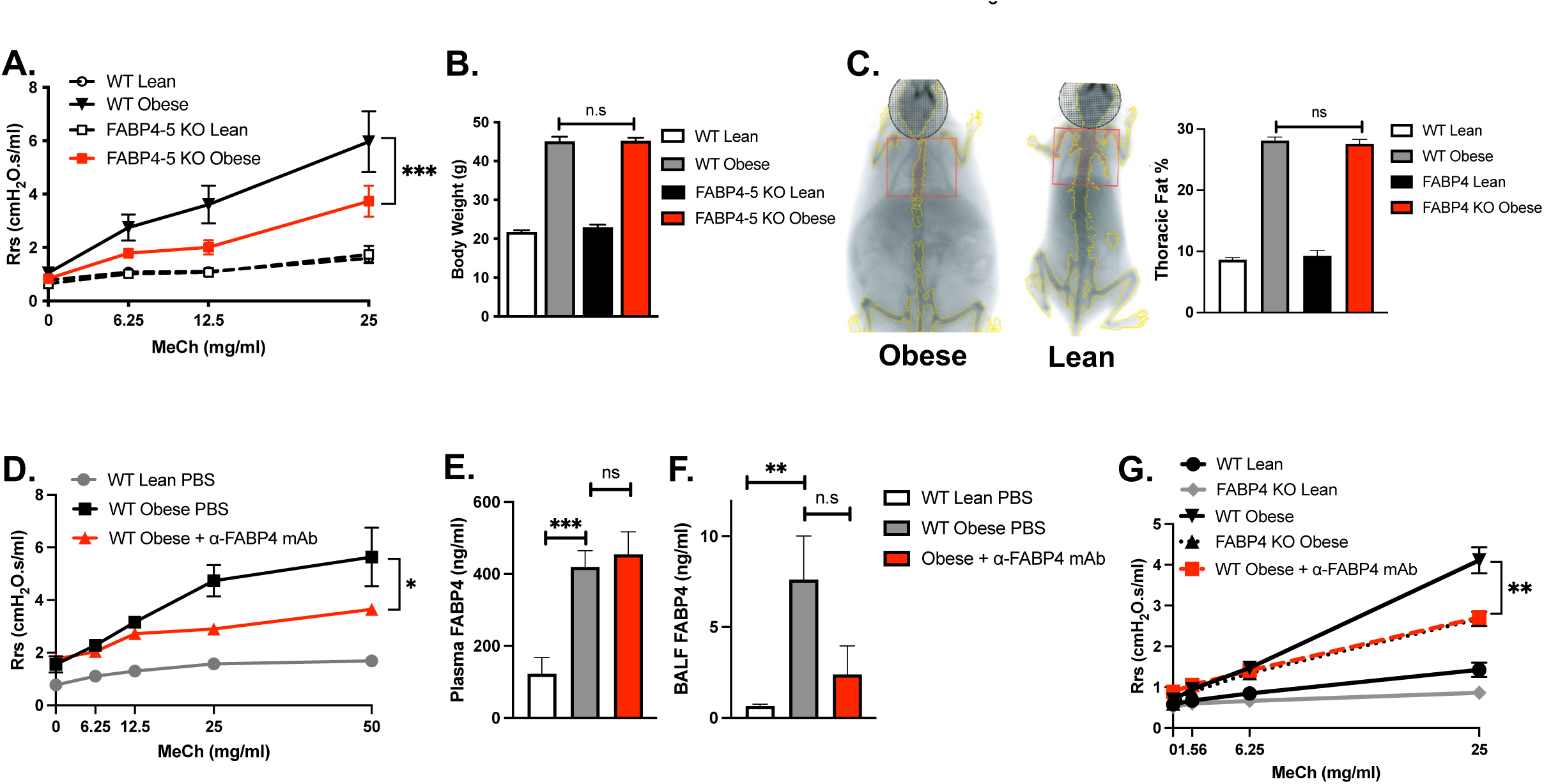
Therapeutic targeting of adipose-derived FABP4 reduces airway hyperresponsiveness in obese mice. **A.** Airway hyperresponsiveness in genetically obese (ob/ob) and lean wild type (WT) and FABP4 and FABP5 double knockout mice (FABP4-5 KO) on a Balb/C background. Measurements were performed using the FlexiVent Fx system with the forced oscillation technique, starting at baseline and then with increasing doses of methacholine (MeCh) up to 25 mg. **B.** Body weights from mice used in panel A (n=4-9 per group). **C.** Representative DEXA scanner body composition images from obese and lean WT mice, and quantification of thoracic fat percentage in WT and FABP4 KO lean and obese (ob/ob) mice (n=3-9 per group). **D.** Measurement of airway hyperresponsiveness in WT lean treated with PBS (WT PBS), WT obese treated with PBS (ob/ob), WT obese (ob/ob) treated with anti-FABP4 monoclonal antibody (mAb) (n=3-8 per group). **E.** Plasma and **F.** BALF FABP4 levels from mice in panel D, measured by ELISA. **G.** Measurement of airway hyperresponsiveness in WT lean, FABP4 KO lean, WT obese (ob/ob), FABP4 KO obese (ob/ob), and WT obese (ob/ob) mice treated with anti-FABP4 monoclonal antibody (mAb) (n=4-7 per group).

In mice and humans, there is increased adiposity in the chest cavity (*14, 55*), which led to the speculation that this may have a role in asthmatic phenotypes of obese subjects. However, this would reflect more of a restrictive lung disease rather than an obstructive lung disease such as asthma. To better understand the degree of thoracic cavity fat deposition in our ob/ob mice, we performed DEXA (dual-energy x-ray absorptiometry) scanning. We compared the adiposity of the genotypes by setting the ROI (region of interest) to their chest. While lean mice had less than 10% fat in their chest cavity, obese mice had almost 30% fat, regardless of the genotype (Fig. 5C). The similar fat percentages in WT ob/ob and FABP4 KO ob/ob mice rules out a purely mechanical effect and instead, suggests a FABP4 protein-dependent impact on the pulmonary dysfunction phenotype in obesity.

Since we showed that obesity increases BALF FABP4 levels, and the absence of FABP4 protects from obesity-driven airway hyperresponsiveness, we decided to explore the therapeutic implication of eliminating the effects of circulating FABP4. We hypothesized that targeting the adipo-pulmonary axis by neutralizing circulating FABP4 may be a potential path to treat obesity-related airway disease. We had previously developed a monoclonal antibody (mAb) raised against FABP4 that successfully reverted metabolic disease in mouse models of diabetes (*18, 36*). Therefore, we decided to test whether we can reverse obesity-related hyperresponsiveness utilizing this monoclonal antibody. We subcutaneously injected ob/ob mice with anti-FABP4 mAb at 30 mg/kg or PBS as a vehicle every fourth day for three weeks (Fig. S5C), followed by measurement of airway hyperresponsiveness. Antibody treatment markedly reduced obesity-induced airway resistance (Fig. 5D). We measured plasma and BALF FABP4, and as expected in both compartments, levels were significantly higher in obesity. While plasma FABP4 levels remained elevated post-antibody treatment (Fig. 5E), BALF levels in antibody-treated mice showed a decreasing trend, though this was not statistically significant (Fig. 5F). To compare the magnitude of the effect of the antibody treatment with genetic deficiency of FABP4, we set up another cohort of mice from a different colony, using littermate FABP4 KO and WT lean and ob/ob mice. We used the same treatment regimen and experimental protocol. In the MeCh challenge test, consistent with our previous findings, ob/ob mice had the highest airway resistance, whereas anti-FABP4 mAb treatment remarkably improved the response, to the same degree as whole-body FABP4 deficiency (Fig. 5G). The antibody treatment did not affect the body weights of the mice in any of the cohorts, again suggesting weight-independent effects of FABP4-neutralization or deficiency on airway function (Fig. S5D and E).

These collective findings underscore the pivotal role of adipose-derived FABP4 in mediating obesity-related airway disease. The consistent protective effects observed through both genetic deletion of FABP4 and neutralization via monoclonal antibodies highlight the critical importance of targeting the adipo-pulmonary axis. By neutralizing FABP4, we unveil a promising therapeutic strategy that holds the potential to transform the treatment landscape for patients suffering from obesity-related airway disease, especially those unresponsive to conventional therapies. This innovative approach paves the way for novel interventions that could significantly improve clinical outcomes and quality of life for this patient population.

## Discussion

Obesity is not an isolated condition; it typically manifests as a conglomerate of interconnected immunometabolic diseases, including airway dysfunction and asthma. Similar immunometabolic underpinnings are evident in non-alcoholic fatty liver, atherosclerotic plaque, and diabetic vasculature, sharing some common molecular etiology (*56, 57*) and may therefore benefit from shared therapeutic interventions, as seen in GLP-1 agonism (*58–61*). Intriguingly, our research identified a similar immunometabolic stress-related signature in obese lungs, highlighting its dependency on FABP4. Our GSEA analysis illustrated that obesity not only increases the risk of asthma but also primes the lung for multiple pathologies, including, but not limited to, infection, cancer, fibrosis, hypercoagulability, acute lung injury, and pulmonary hypertension. For example, the top enriched pathway in obese lungs was neoplastic transformation (Fig. S1C) and GSEA revealed pulmonary vascular endothelium damage with dramatic upregulation of the Vegfa gene (Fig. S2D), which is known to be upregulated by FABP4 (*43*).

Our comprehensive study focused on detailed definition of obesity-related airway disease by characterizing the lungs of obese mice through various analyses. These included histology, imaging, single-cell RNA sequencing, BALF proteomics, and lipid profiling. Notably, we examined airway physiology in genetic and diet-induced obesity models in a naïve state, aiming to decipher the multiple cellular clusters and merging pathways that directly contribute to airway disease in obesity. One of our key mechanistic discoveries was identifying lipotoxic stress-induced disruptions in lipid metabolism, which led to maladaptive surfactant metabolism in obesity. This alteration involved downregulation of genes involved in lipid biosynthesis and modification, particularly Scd1, that would impact surfactant metabolism. Interestingly, our findings mirrored previous studies linking Scd1 downregulation to decreased surfactant protein and asthma pathogenesis in mice and humans (*62*). Supporting this literature, we also found dramatically lower surfactant proteins in obesity that were partly rescued by FABP4 deficiency. This finding is exciting and highly relevant to asthma, as it aligns with the established pathology in humans, and successful use of surfactant therapeutic interventions in animal asthma models (*50, 51, 63, 64*). However, the feasibility of surfactant treatment in humans is challenging and requires intubation. Therefore, its use is limited to newborns with severe respiratory distress syndrome (RDS), whereas neutralizing hormonal FABP4 might be a viable and broadly available alternative.

We previously showed that FABP4-deficient mice are also protected from allergen-induced asthma (*38*). In that study, bone marrow transplantation experiments led us to the conclusion that parenchymal cells were the driver of airway disease. After discovering the nutrient-regulated presence of FABP4 in circulation (24, 33, 65), as well as in BALF in both mice and humans in this study, we hypothesize that circulating FABP4 could mediate airway disease associated with obesity by influencing cellular behavior specifically within the bronchoalveolar space, and more broadly, throughout the lung. In this study, despite comparable total body masses and thoracic fat percentages across various models and backgrounds, we provide evidence that obesity-related pathologies might arise not merely from increased fat mass. Rather, signaling molecules, adipokines, and inflammatory molecules emanating from different cellular components determine the disease pathogenesis. This includes the airways, establishing an intriguing pathological adipo-pulmonary axis.

We have now identified the presence of FABP4 in the bronchoalveolar space, regulated with obesity in both mice and humans. Remarkably, its regulation in humans was more pronounced in women with obesity, where airway disease incidence is significantly higher, implicating FABP4 in the disease’s pathogenesis. Our transplantation and implantation experiments pinpointed adipose tissue as the source of FABP4 in the airways (Fig. S4J). As a limitation, we cannot differentiate the relative impact of different fat depots or determine whether intrapulmonary fat cells directly affect FABP4 exposure to the airways. Importantly, in obesity-related airway disease characterized by increased airway resistance during bronchoprovocation, significant reversal was observed with either FABP4 deficiency or monoclonal antibody targeting of circulating FABP4. These results underscore FABP4’s critical role in driving airway pathology via the adipo-pulmonary link and highlight therapeutic avenues for targeted intervention.

Human sample analysis from the Nurses’ Health Study and Health Professionals Follow-up Study (*65*) offered valuable insights, albeit with a lower asthma frequency, underestimating FABP4’s impact on asthma. Larger asthma-focused cohorts are imperative to strengthen our findings’ statistical power. Furthermore, exploring FABP4’s role in the relationship between BMI and asthma severity in humans remains intriguing. Given the complex and evolving biology of the intracellular versus circulating forms, we should carefully interpret FABP4’s role in different asthma subtypes. Our study supports and translates the role of circulating FABP4 in non-eosinophilic, non-neutrophilic obesity-related airway-disease prevalent in female patients. We cannot rule out the possible differential role of intracellular FABP4 in macrophages or eosinophils in other subtypes of asthma.

By elucidating the role of a fatty acid-binding protein, our study contributes significantly to the understanding of obese asthma phenotype and unveils the intricate pathological connection between obesity and airway disease. Adipose-born FABP4, higher in obesity, regulates distant organs’ immune and metabolic functions through its pleiotropic effects (*17*). Our new data establishes an unexpected and dramatic impact of FABP4 in the lungs and airways. The impact of this work can be summarized in three pillars. First, it puts forward a concept that a novel adipokine can target and regulate the airways. Second, this adipo-pulmonary axis may serve as a marker of obesity-related airway disease. Third, it provides an exciting opportunity for translation of the therapeutic utility of neutralizing a circulating adipokine to prevent and treat obesity-related asthma. Targeting the adipo-pulmonary axis via FABP4 inhibition might be a novel treatment that addresses the unmet medical need for the management of obesity-related asthma.

## Materials and Methods

### Study Design

The objective of the study was to investigate the effects of obesity on lung and airway physiology, define the obesity-related asthma phenotype, and explore the role of circulating FABP4 in this context. We also tested therapeutic properties of a monoclonal antibody raised against FABP4 on airway hyperresponsiveness in obese mice. We validated the presence of FABP4 in human samples obtained from cohorts collected at Asthma Research Center in Brigham and Women’s Hospital, Boston, MA. *In vivo* results shown for each experiment were replicated in at least two separate cohorts and in different backgrounds of mice including C57BL/6 in-house, C57BL/6J, and BALB/c in-house. Histological sections were quantified blinded in the lab from sections stained and scanned at Histowiz (Long Island City, NY). Single cell RNA sequencing was performed at the Genomics and Cell Biology core at NORCH (Nutrition Obesity Research Center at Harvard) from the cell suspensions prepared in the lab. Proteomics studies were performed at the Harvard Chan Multi-Omics Platform (ChAMP) in the Department of Molecular Metabolism from bronchoalveolar lavage samples collected in the lab. *In vitro* constructs of 3T3-L1 mouse preadipocytes expressing NanoLuc-tagged FABP4 were prepared and validated in the lab. Lung imaging by computed tomography was performed in the lab using micro-CT (eXplore CT 120 micro-CT, GE Healthcare, Waukesha, WI). *In vivo* luminescence images were captured using an IVIS Spectrum *in vivo* imaging system (Perkin Elmer, MA). Airway physiology experiments were performed in the lab using a flexiVent FX system (SCIREQ, Canada). In all *in vivo* experiments the elimination criteria for outliers were based on the visible health of the individual mice, known experimental/technical errors or findings more than 2 SDs from the mean.

### Animals

Animal care and experimental procedures were performed with the approval from the Harvard Medical Area Standing Committee on Animals. All mice were kept on a 12-hour light/dark cycle. Female leptin-deficient (ob/ob - Strain #:000632) and wild type (lean - Strain #000664) control mice on C57BL/6J background were purchased from the Jackson Laboratory. Cohorts for diet-induced obese mice and regular diet control mice were set in-house and fed 60% kcal fat (Research Diets Inc., D12492i) or minimum 4.5 % kcal fat diet (5053 - PicoLab® Mouse Diet 20) starting at 4 weeks of age for 8 weeks. BALB/c ob/ob and BALB/c *Fabp4*-knockout ob/ob mice and their controls were kept on regular chow diet (5053 - PicoLab® Mouse Diet 20). All *in vivo* experiments were performed on mice between 10-12 weeks of age unless otherwise stated. Only female mice were used for the airway physiology and single cell RNA sequencing experiments. In-house *Fabp4*-KO ob/ob on C57BL/6 background were backcrossed to BALB/c for over 9 generations.

### Human Samples

All the human samples analyzed in this study were received in a de-identified manner. Serum samples of human asthmatic and non-asthmatic subjects were from The Nurses’ Health Study (121,700 women) and Health Professional’s Follow-up Study (51,529 men) which are ongoing surveys tracking chronic diseases. Samples used in this study were randomly selected and were previously analyzed for cardiovascular outcomes (unpublished data). The Institutional Review Board of the Brigham and Women’s Hospital and the Harvard School of Public Health Human Subjects Committee approved the study protocol. Human BALF samples were provided anonymously by the Asthma Research Center at Brigham and Women’s Hospital and were collected during elective bronchoscopies with informed consent. All the BALF sample collection was performed with the approval of the Mass General Brigham Institutional Review Board (IRB approval #s: 2012P001426, 2012P002190, 2010P000170, 2012P001528). All the available samples were collected according to the established standardized protocol by the Severe Asthma Research Program(*66*).

### Plasma and BALF FABP4 Measurements

Plasma and BALF FABP4 measurements were done using an in-house sandwich ELISA. Antibodies used (clone 351.4.2E12.H1.F12 for capture, HRP-tagged clone 351.4.5E1.H3 for detection) were produced for the Hotamışlıgil lab at Dana-Farber Cancer Institute Monoclonal Antibody Core, Boston, MA (*36*). Human recombinant FABP4 (R&D Systems) was used as a standard. Lower limit of detection was 0.0625 ng/ml. Human serum sample FABP4 measurements were done with a commercial ELISA kit (Biovender), according to the manufacturer’s recommendations. All samples were measured in duplicates.

### Lung Single Cell Suspension Preparation

Mice were anesthetized with xylazine (7.2 mg/kg ip, XylaMed, Bimeda Inc.) followed by pentobarbital (70 mg/kg ip, Nembutal Sodium, Oak Pharmaceuticals), 5 minutes apart. Once the mice reached a surgical plane of anesthesia, e.g. non-reactive to toe pinch, the chest cavity was opened, and the lungs were perfused with 20 ml isotonic saline through the right ventricle. The lobes were removed and placed in a petri dish with cold DMEM with 10% FBS and kept on ice until all the lungs were harvested. The lung of each mouse was put into a 50 ml conical tube containing 5 ml DMEM with 2% FBS and cut into 2-3 mm pieces using scissors and kept on ice. Once all the lungs were cut, 5 ml of warm 2X digestion buffer (Liberase TL 0.4 mg/ml, elastase 0.1 mg/ml, DNase1 0.1 mg/ml in DMEM with 2% FBS) was added to each tube and incubated in a shaker at 37°C for 30 minutes. Following the incubation, the tissue clump was dislodged by pipetting up and down multiple times with a 10 ml serological pipette and then filtered through a 100 µm cell strainer into a clean 50 ml conical tube and kept on ice. The undigested tissue was removed from the strainer, placed into a clean 50 ml tube with 5 ml of 1X digestion buffer and incubated in a shaker at 37 °C for 20 minutes. The digest was then strained through a 70 µm cell strainer into the 50 ml tube on ice from the previous round. After this second round of filtration, an equal volume of cold DMEM with 2% FBS was added, and the cells were centrifuged at 1500 rpm at 4°C for 10 minutes. The supernatant was discarded, and the pellet was resuspended in 5 ml of red blood cell lysis buffer (Thermo Fisher) and incubated at room temperature for 5 minutes, shaking every minute. The cells were centrifuged at 1500 rpm for 10 mins at 4°C and the pellet was resuspended in 5 ml single-cell suspension buffer (2 mM EDTA, 0.5 BSA, 1% FBS in DPBS) and the cells were counted (Bio-Rad TC20) to prepare 1×10^6^ aliquots.

### Single-Cell RNA-Sequencing and Data Analysis

Single-cell RNA-seq library construction was performed on a Chromium 10x instrument using Chromium single cell 3’ reagent v3.0 kit (10x Genomics), followed by sequencing on an Illumina NextSeq 2000 instrument, which resulted in approximately 100-150 million reads per sample. Initial processing of scRNA-Seq data was performed using CellRanger software (v6.0.0) (*67*). The reads were aligned to the mm10 mouse reference genome, with a mapping rate of ∼90%, followed by the calculation of read counts per gene in each cell. Further analysis was performed using Seurat 3.2.3 (*68*). Cells with <200 expressed genes and genes expressed in <3 cells were filtered out, followed by the exclusion of cells with > 10% of mitochondrial transcripts. Read counts for each sample were normalized using the NormalizeData function in Seurat. Using the FindVariableFeatures function, 2000 features were selected to be used in Principal Component Analysis (PCA). All individual samples were integrated using the Seurat canonical correlation analysis (CCA) method. Integration anchors were determined using the FindIntegrationAnchors function and then used in IntegrateData function. UMAP plots and cell clusters were generated using the RunUMAP and FindClusters functions, followed by manual cell type annotation based on type-specific gene markers for each cluster. In addition, cell types corresponding to individual cell clusters were inferred using ELeFHAnt (*69*) by the annotation transfer from the mouse lung atlas (*70*). Feature plots for individual genes were generated using the FeaturePlot function of Seurat. Differentially expressed genes between conditions within cell populations of interest were identified using the FindMarkers function. Dot plots of individual gene expression were generated using the DotPlot function. Pathway enrichment analyses were performed using EnrichRbased on Human WikiPathway 2021 database and using GSEA based on the MSigDB databases and curated gene lists for functional categories of interest.

### LC-MS/MS Proteomics Analysis

Lavage fluid from the lungs of mice was buffered with 200 mM HEPES (4-(2-hydroxyethyl)-1-piperazineethanesulfonic acid) pH 7.5. Disulfide bonds were reduced using 5 mM dithiothreitol (Sigma-Aldrich) at 37°C for 1 h, followed by alkylation of cysteine residues using 15 mM iodoacetamide (Sigma-Aldrich) in the dark at room temperature for 1 h. Excessive iodoacetamide was quenched using 10 mM dithiothreitol. Proteins were precipitated overnight at 4°C using ice-cold acetone/methanol mixture (9:1 v/v). Samples were centrifuged at 6,000 x g for 75 min at 4°C. Protein pellets were washed twice using 1 ml ice-cold methanol. Protein pellets were dissolved using 200 mM HEPES pH 7.5 prior to digestion using sequencing grade trypsin (Worthington Biochemical) at 37°C for 16 h. Digested peptides were subsequently desalted using self-packed C18 STAGE tips (3M Empore^TM^) (Rappsilber, 2003) for LC-MS/MS analysis. Desalted peptides were resolubilized in 0.1% (v/v) formic acid and loaded onto HPLC-MS/MS system for analysis on an Orbitrap Q-Exactive Exploris 480 (Thermo Fisher Scientific) mass spectrometer coupled to an FAIMS Pro Interface system and EASY-nLC 1000 (Thermo Fisher Scientific) with a flow rate of 300 nl/min. The stationary phase buffer was 0.1 % formic acid, and the mobile phase buffer was 0.1 % (v/v) formic acid in 80% (v/v) acetonitrile. Chromatography for peptide separation was performed using an increasing organic proportion of acetonitrile (5 – 40 % (v/v) over a 120 min gradient) on a self-packed analytical column using a PicoTip^TM^ emitter (New Objective, Woburn, MA) with Reprosil Gold 120 C-18, 1.9 µm particle size resin (Dr. Maisch, Ammerbuch-Entringen, Germany). The mass spectrometry analyzer operated in data independent acquisition mode at a mass range of 300–2000 Da, compensation voltages of −50/−70 CVs with survey scan of 120,000 and 15,000 resolutions at the MS1 and MS2 levels, respectively.

Mass spectrometry data were processed by Spectronaut^TM^ software version 15 (Biognosys AG) (*71*) using directDIA^TM^ analysis with default settings, including: oxidized methionine residues, biotinylation, protein N-terminal acetylation as variable modification, cysteine carbamidomethylation as fixed modification, initial mass tolerance of MS1 and MS2 of 15 ppm. Protease specificity was set to trypsin with up to 2 missed cleavages allowed. Only peptides longer than seven amino acids were analyzed, and the minimal ratio count to quantify a protein was 2 (proteome only). The false discovery rate (FDR) was set to 1% for peptide and protein identifications. Database searches were performed using the UniProt *Mus musculus* strain database containing 54,822 entries (Dec 2020). High precision iRT calibration was used for samples processed using the published nanospray conditions (*72*). Protein tables were filtered to eliminate identifications from the reverse database and common contaminants. The mass spectrometry proteomics data have been deposited to the ProteomeXchange Consortium via the PRIDE (*73*) partner repository with the dataset identifier PXD051131.

### FlexiVent FX Measurements to Evaluate Airway Function

Mice were anesthetized with xylazine (7.2 mg/kg ip) followed by pentobarbital (70 mg/kg ip), 5 minutes apart. Once a surgical level of anesthesia was reached, i.e. non-reactive to toe pinch and having a regular breathing pattern, the mice were placed on a surgery board in supine position. The neck was fixed on the board in a hyperextension position by hooking the incisors with an elastic band anchored to the board, and the throat area was cleaned with 70% ethanol. A midline incision to the lower neck was made below the mandible. The submaxillary glands and the muscle layer were gently separated to expose the trachea. After cutting the membrane surrounding the trachea, a suture was passed underneath it using a forceps. A small incision was made between two cartilage rings near the larynx, and an 18-gauge metal cannula (resistance: 0.36 – 0.40 cmH2O·s/ml) was inserted caudally at a distance of 4-5 rings and secured with the previously placed suture. Once the mouse was tracheotomized, the cannula was connected to the flexiVent FX system with nebulizer and two deep inflation maneuvers were performed 30 seconds apart to recruit all the airways and standardize the lung volume. The mechanical ventilation was set at 150 breaths/min with tidal volume of 10 ml/kg and positive end-expiratory pressure (PEEP) of 3 cmH_2_O. To induce muscle relaxation, pancuronium (0.72 mg/kg, Sigma P1918) was administered, and the mice were ventilated for 5 minutes for the drug to take effect. At the end of the 5 minutes, spontaneous inspiratory efforts were checked by stepwise pressure-volume curves. Baseline respiratory mechanics and airway resistance to methacholine challenge were measured by a sequence composed of two deep inflations (6 sec, 3-30 cmH_2_O), 12 SnapShot-150 (1.2 s, 2.5 Hz), and 12 QuickPrime-3 (3 sec, 1-20.5 Hz) maneuvers.

### FABP4-NanoLuc-Expressing Mouse 3T3-L1 Cell Line Preparation

#### Cloning

Nucleotides encoding human FABP4-NanoLuc was synthesized by Integrated DNA Technologies (IDT Inc, Coralville, Iowa). The nucleotides were cloned into the entry vector, pENTR1A no ccDB, at EcoRI/NotI sites and then the destination vector, pLenti CMV Puro DEST (w118-1), using Gateway™ LR Clonase™ II Enzyme mix (ThermoFisher #11791020).

#### Stable cell

HEK293T cells (ATCC CRL-3216) were transfected with pLenti-hFABP4-NanoLuc, packaging plasmid psPAX2, and enveloping plasmid pMD2.G using Lipofectamine LTX plus (ThermoFisher #15338100). Forty-eight hours after transfection, media containing the lentiviral particles was filtered through 0.45 μm syringe filter and added onto 3T3-L1 mouse preadipocytes (Zenbio SP-L1-F). Forty-eight hours after infection, the cells were treated with 2mg/ml puromycin for 2 weeks for stable cell selection.

#### Differentiation

hFABP4-NanoLuc-3T3-L1 preadipocytes cultured on 10-cm plates to 100% confluence were differentiated into mature adipocytes using a previously established protocol (*74*). Briefly, the cells were incubated in differentiation media (DMEM, 10% FBS, 1% penicillin, 1% streptomycin, 5 µg/ml insulin (Sigma I9278), 1 µM dexamethasone (Sigma D4902), 500 µM IBMX (Sigma I5879), and 2 µM rosiglitazone (Cayman 71740)) for 48 hours then switched to maintenance media (DMEM, 10% FBS, 1% penicillin, 1% streptomycin, 5 µg/ml insulin), and changed every 48 hours. Mature adipocytes were collected at day 8-10. Cells were lifted with 0.25% trypsin-EDTA (Gibco 25200056) and spun at 50 x g for 10 minutes. The pellets were resuspended in sterile PBS for injection into mice.

#### NanoLuc assay

*In vitro* NanoLuc luciferase activity was measured using the Nano-Glo® Luciferase Assay System from Promega (#N1110) following the manufacturer’s protocol. Briefly, the assay reagent was prepared by diluting the substrate (furimazine) into the assay buffer at 1:100. In a 96-well half-area white flat-bottom microplate (Corning 3883), 10 µl sample, 40 µl PBS, and 50 µl assay reagent were mixed and incubated at room temperature for 10 minutes. The luminescence was read on a SpectraMax® Paradigm® Multi-Mode Microplate Reader (Molecular Devices, San Jose, CA).

#### List of plasmids used

- pENTR1A no ccDB (w48-1) was a gift from Eric Campeau & Paul Kaufman (Addgene plasmid # 17398; http://n2t.net/addgene:17398; RRID: Addgene_17398)
- pLenti CMV Puro DEST (w118-1) was a gift from Eric Campeau & Paul Kaufman (Addgene plasmid # 17452; http://n2t.net/addgene:17452; RRID: Addgene_17452)
- psPAX2 was a gift from Didier Trono (Addgene plasmid # 12260; http://n2t.net/addgene:12260; RRID: Addgene_12260)

### Adipose Tissue Transplantation Experiments

Adipose tissue transplantation experiments were performed using WT and FABP4 KO lean and obese mice on BALB/c background. WT ob/ob mice were the donors, and lean FABP4 KO mice were the recipients in our transplantation experiments. Briefly, both the donor and the recipient mice were anesthetized simultaneously with 100 mg/kg ketamine / 10 mg/kg xylazine. Once the donor mice reached the surgical level of anesthesia, the abdomen was shaved and disinfected. A sterile drape was then placed on the mice with an opening to expose the mid-abdomen. The skin and muscle layers were opened vertically and the left perigonadal white adipose tissue was exposed and excised, then placed into a sterile DPBS-containing petri dish. The donor was then euthanized while under anesthesia. The excised tissue was weighed, and 2 ± 0.2 grams of adipose tissue was cut into small pieces. The abdominal skin of the anesthetized recipient mice was shaved and disinfected, and then the skin and muscle layers were cut vertically along the midline from the mid abdomen to 0.5-1 cm above the bladder. Half of the adipose tissue pieces were placed on the left and the other half on the right side of the peritoneal cavity. The transplanted tissue pieces were kept as small as possible, as larger tissues tend to be more antigenic and are more likely to be rejected and/or become necrotic. The peritoneal muscle layer was closed with absorbable 5-0 sutures, and suture clips were used to close the abdominal skin. For pain management, Buprenorphine ER was injected immediately after surgery (0.5mg/kg subcutaneously). Mice were individually placed in clean cages on top of a 37°C circulating water pad until they recovered from anesthesia. The mice were monitored daily for signs of infection and acute rejection-related systemic illness over a four-day period.

### Adipocyte Cell Line Implantation Experiments

3T3-L1 cells were implanted on day 8-10 of differentiation (complete differentiation takes 10-14 days). Cells were resuspended in sterile PBS (5 x 10^6 cells per 100 µl) and kept on ice until implantation. Depending on the experiment, 5 to 60 x 10^6 cells were implanted by subcutaneous injection using a syringe with a 22G needle into the left lower back of the mice. Following a 24-hour post-injection period, 50 µl of blood was collected from the tail vein daily for two days. On the third day, a final blood sample was collected, and the mice were euthanized to collect BALF for FABP4 measurements and luciferase assays.

### FABP4 Secretion Experiments

Secretion experiments were performed by administering lipolytic agents. For *in vivo* experiments pan-beta agonist isoproterenol (Tocris Biosciences) in PBS was administered at a dose of 10 mg/kg intraperitoneally (ip). Blood samples were collected from the tail vein at baseline (0), 45-, 60-, 90-, and 180-minutes following injection, depending on the experimental design. Plasma samples were used for FABP4 measurements. For *in vitro* experiments, differentiated Nluc-tagged FABP4-expressing 3T3-L1 adipocytes were treated with forskolin (FSK, 20μM) or Isobutylmethylxanthine (IBMX, 1mM) for 1 hour and the conditioned media was collected for FABP4 measurements by ELISA.

### IVIS Imaging

An IVIS^®^ Spectrum (Perkin Elmer) *in vivo* bioluminescence imaging system was used both for live animal and *ex vivo* organ imaging. For the *in vivo* experiments, mice were anesthetized with 1-2% isoflurane and baseline images were taken prior to substrate injection. The mice were then injected with 100 µl substrate (40x diluted from the stock) Nano-Glo® Fluorofurimazine *In Vivo* Substrate (FFz, Promega, Madison, WI) via the tail vein (*75*). The mice images presented in the manuscript were imaged 30 minutes after systemic substrate injection. Of note: As the peak luciferase signal intensity changes over time, adjustments may be necessary if imaging occurs beyond 3 days post-transplant. Ideally, images should be taken within 10-30 minutes after substrate injection in these cases. In the experiment for organ imaging, after a small incision to the atrium, the mice were perfused with 20 ml saline through the left ventricle 30 minutes after systemic substrate injection. Subsequently, the organs (lung, spleen, liver, heart, kidney, and perigonadal adipose tissue) were imaged. In a separate experiment, following perfusion, the lungs were flushed via lung lavage through the trachea with 500 µl PBS, repeated 10 times to clear the bronchoalveolar space before imaging.

### MH-S Cell line Cytokine Secretion Experiments

The mouse alveolar macrophage cell line MH-S (ATCC# CRL-2019) was cultured in growth medium (RPMI-1640 supplemented with 10% FBS and 50 mM 2-mercaptoethanol). Cells were seeded at 1.5 x 10^5 cells / well in 12-well plates and cultured to confluency. For the experiment, the cells were washed twice with PBS and then treated with recombinant FABP4 (100 ng/ml), TNF-a (10 ng/ml), or left untreated (vehicle-treated with PBS, no drug or protein) in 1 ml starvation medium (RPMI-1640 + 0.1% BSA) for 18 hours. Next day, the cells were treated in growth medium for 8h. After 8 hours, the conditioned media was collected, and the cells were washed twice with PBS and immediately frozen in liquid nitrogen until further processing. Separate sets of cells were also collected and frozen in RIPA buffer containing 2 mM activated sodium orthovanadate (Na_3_VO_4_) (New England Biolabs) and 1% protease inhibitor cocktail (MilliporeSigma) for protein isolation. All treatments were done in triplicates. For proteome profiling (R&D Proteome Profiler Mouse Cytokine XL, cat# ARY028), frozen conditioned media was thawed overnight at 4 °C and then processed according to the manufacturer’s instructions. Images of the membranes were captured using a ChemiDoc MP Imaging System (Bio-Rad Laboratories). Protein signal quantification was performed using Fiji/ImageJ2 (version 2.3.0) image processing software with a pre-made automated ROI template.

### Body Composition using DEXA and CT Scanning

Body composition was measured using a DEXA scanner (Lunar PIXImus2, GE Healthcare, Waukesha, WI) in anesthetized mice (100 mg/kg ketamine / 10 mg/kg xylazine, ip). After obtaining the whole-body scan image, a standardized ROI was placed on the thorax region (separate for lean and obese mice) to measure the amount and percentage of fat in this area. Whole mouse tomography images with a focus on the thoracic region were obtained using a CT scanner (eXplore CT 120 micro-CT, GE Healthcare, Waukesha, WI), as previously described (76). CT scanning was performed as a terminal experiment right after euthanasia, and images were obtained at maximum inspiration by inflating the lungs prior to imaging. For obese mice, tidal volume (10 ml/kg) for the ideal weight (weight of the lean litter mate) (*76*) was calculated, and 30% additional volume (10 ml/kg x 1.3 = maximum inspiration) was added. The air volume was delivered by a 10mL syringe attached to a tracheostomy cannula. The image of the representative WT obese mouse was reconstructed from the exported DICOM (Digital Imaging and Communication in Medicine) images. DICOM images were converted to JPEG format using Horos ^TM^ software to generate the figure. Horos is a free and open-source code software (FOSS) program that is distributed free of charge under the LGPL license at Horosproject.org and sponsored by Nimble Co LLC d/b/a Purview in Annapolis, MD USA.

### Histology, Immunohistochemistry, and Intrapulmonary Fat Measurements

The left lung was used for histological examination. Following euthanasia of the mouse, the chest was opened, and the right lung was ligated at the right main bronchus, harvested, and promptly frozen for further analysis. Subsequently, 500 µl of 10% zinc formalin was instilled into the left lung via the trachea. The lung was then ligated at the left main bronchus level, harvested, placed in a cassette, and immersed in a jar containing 10% zinc formalin. After 24 hours of fixation, the tissue was rinsed under running tap water for 5 minutes and stored in 70% ethanol. Paraffin embedding, sectioning, H&E staining, and immunohistochemistry (IHC) were carried out by HistoWiz, Inc. (Long Island City, NY), an automated histology platform. The anti-FABP4 antibody (Abcam, #13979) was used to stain for FABP4, while the anti-perilipin antibody (Cell Signaling Technology, #9349) was used for intrapulmonary fat cell detection. Measurements of fatty areas were performed blindly. Fat cell quantification was done using QuPath software v0.5.0 (77). Initially, the entire surface area of the left lobe was computed by outlining the pleura. Fat cells were then identified, and their total surface area was quantified. Using these measurements, the fat percentage relative to the total surface area was calculated. The histological slides used in this analysis included representations of all major and minor airways and vasculature.

### Statistical Analysis

GraphPad Prism software (Version 10.2.1) was used for statistical analysis and graphing our results. Data are presented as means ± SEM. Unpaired Student’s t-tests (2 groups), one-way ANOVAs with post-hoc analyses (3+ groups, 1 variable), two-way ANOVAs, and Spearman’s correlation were used. Significance was set at p≤0.05 and indicated as follows: *p<0.05, **p<0.01, and ***p<0.001. p values between 0.05 and 0.1 were considered as indicating a trend.

## Acknowledgements

We would like to thank the members of the Hotamisligil laboratory, past and present, for their helpful and constructive discussions throughout the years. Specifically, we extend our thanks to Jani Saksi and Kacey Prentice for their assistance with tail vein injections and heart perfusion. We are also very thankful to our funding agencies for their generous support, and to the Nutrition Obesity Research Center at Harvard (NORCH) for the core facility utilization and collaboration.

## Author contributions

Conceptualization: MFB, GT, GSH

Methodology: MFB, GT, GSH, ANA, GYLS, KI, KC, RS, ZWL

Investigation: MFB, GT, GSH

Visualization: MFB, GT, ANA, KC, RS

Funding acquisition: MFB, GT, GSH

Supervision: GSH, EI, RS

Writing- original draft: MFB, GT, GSH

Writing- review and editing: MFB, GT, GSH, ANA, GYLS, KI, ND, KC, RS, EI

MFB and GT had equal contributions on all the categories above.

## Data and materials availability

All datasets and raw data generated during the current study are available from the corresponding author on reasonable request.

## Figure Legends

**Supplemental Figure 1.**
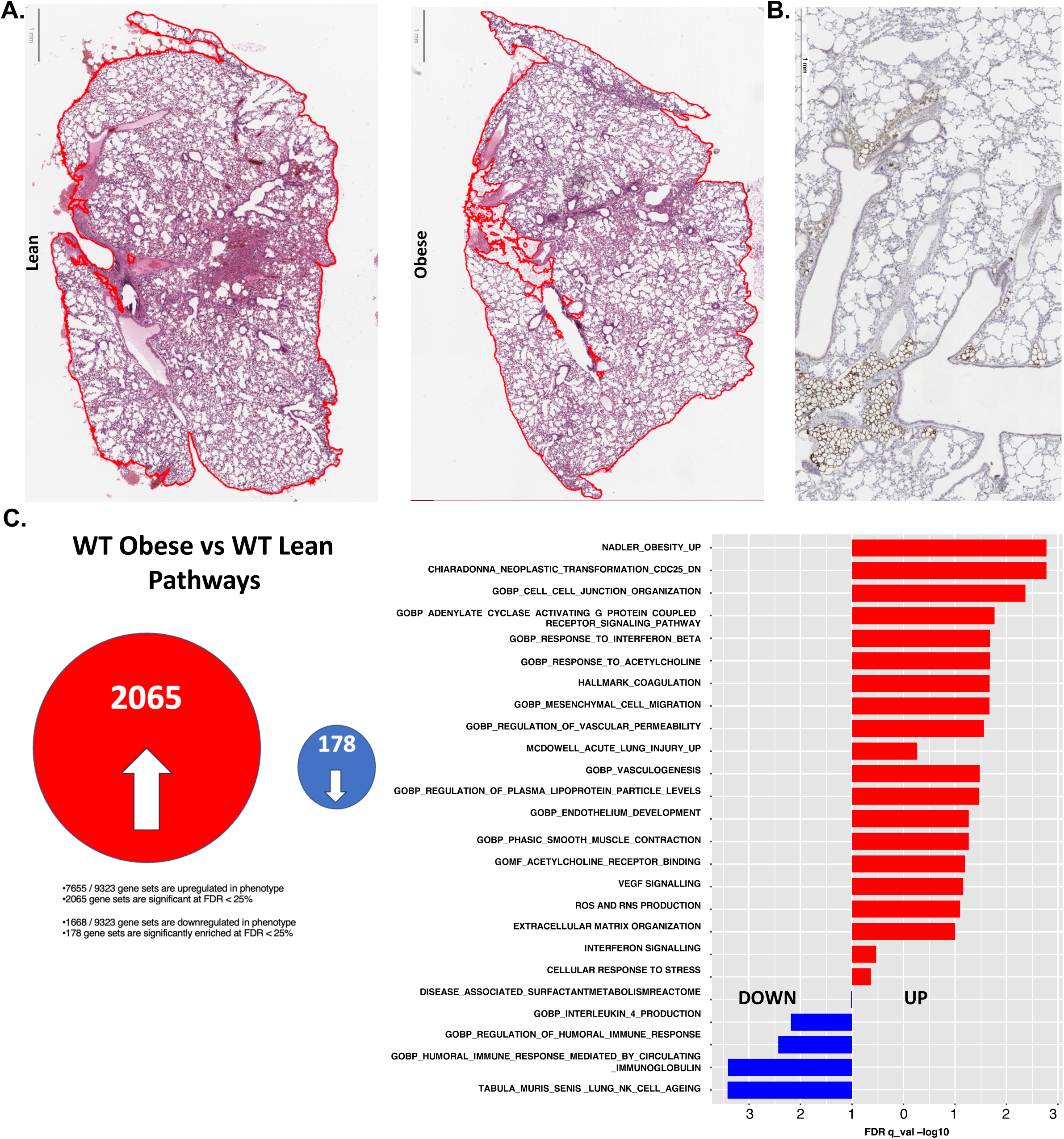
**A.** Representative images of H&E-stained lung sections from lean and obese mice, with marked areas for quantification of fat percentage. **B.** Immunohistochemistry showing perilipin staining of adipocytes in a lung section from an obese mouse. **C.** GSEA analysis of single-cell RNA-seq from whole lungs of WT obese vs. WT lean mice, revealing 2,065 upregulated pathways and 178 downregulated pathways in obese lungs. The right panel shows the top 25 enriched pathways in obese lungs, ranked by FDR (False Discovery Rate) q-value.

**Supplemental Figure 2.**
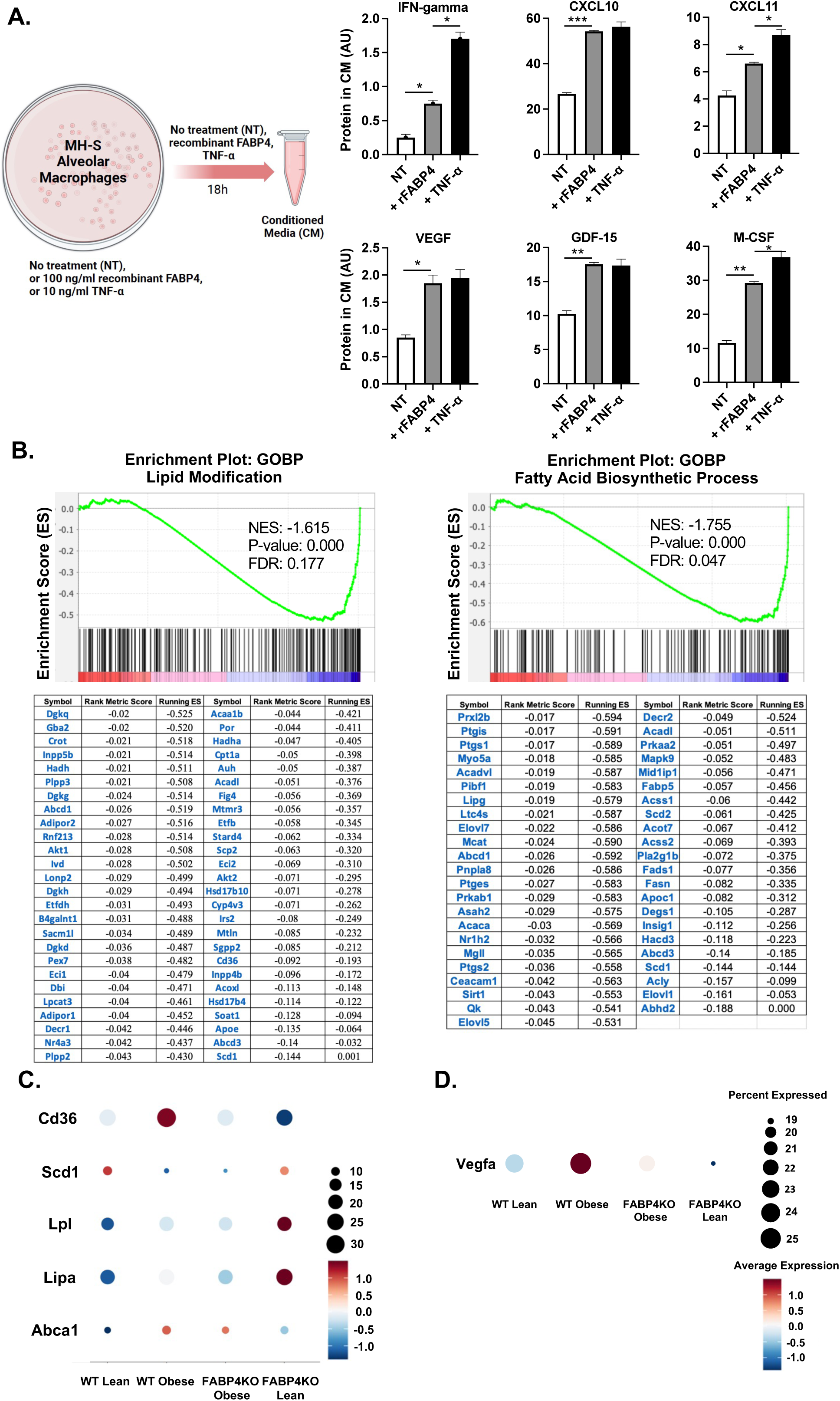
**A.** Left: Cartoon explaining the experimental protocol for measuring secreted proteins in MH-S alveolar macrophages treated with recombinant FABP4, TNF-alpha, or vehicle treatment (NT). Right: Graphs showing protein levels of IFN-gamma, CXCL10, CXCL11, VEGF, M-CSF, and GM-CSF in conditioned medium. The Proteome Profiler Cytokine Array (R&D Systems) was used to measure the secreted proteins. **B.** GSEA enrichment plots and core enriched genes for lipid modification and fatty acid biosynthesis pathways from single-cell preparations of whole lungs in WT obese vs. WT lean mice. **C.** Dot plots showing differential expression of lipid metabolism genes (Cd36, Scd1, Lpl, Lipa, and Abca1) in lungs from WT lean, WT obese, FABP4 KO obese, and FABP4 KO lean mice. **D.** Dot plot of Vegfa gene expression in endothelial cell clusters from lungs of WT lean, WT obese, FABP4 KO obese, and FABP4 KO lean mice.

**Supplemental Figure 3.**
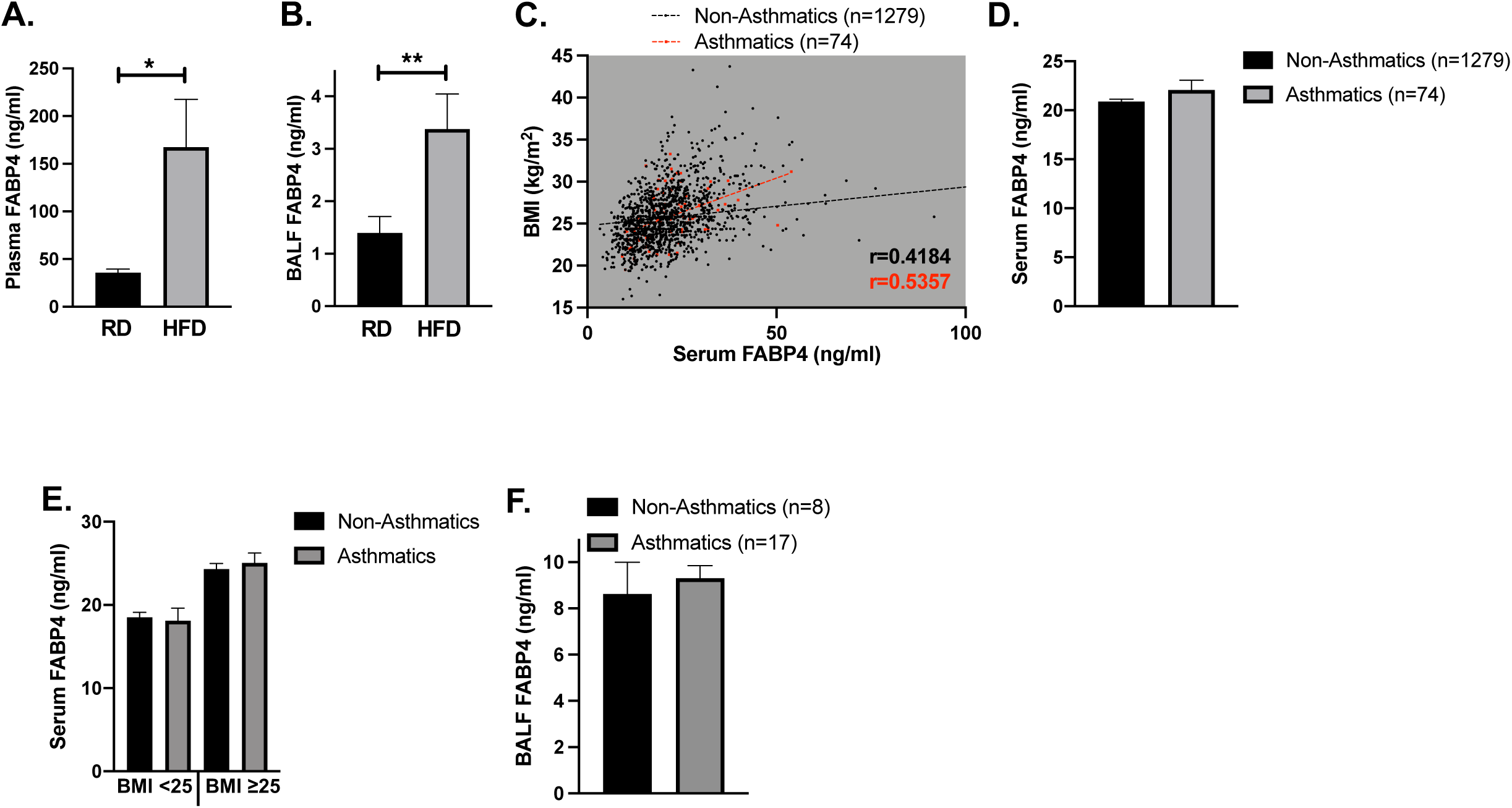
**A.** Plasma FABP4 levels and B. BALF FABP4 levels, measured by ELISA in lean (regular diet, RD) and obese mice (high-fat diet, HFD) (n=16-23 per group). **C.** Correlation of BMI and serum FABP4 levels in males with and without asthma, calculated by Spearman’s correlation test. In non-asthmatics: r=0.4184, p<0.0001. In asthmatics: r=0.5357, p<0.0001. **D.** Serum FABP4 levels in males with and without asthma. **E.** Serum FABP4 levels in males with and without asthma, stratified by lean (BMI < 25) vs. overweight/obese (BMI ≥ 25) status. Sample sizes were 550 and 32 for lean males without asthma and with asthma, respectively. In the overweight/obese groups, sample sizes were 729 and 42 for males without asthma and with asthma, respectively. **F.** BALF FABP4 levels in males with and without asthma, with sample sizes of 17 and 8, respectively.

**Supplemental Figure 4.**
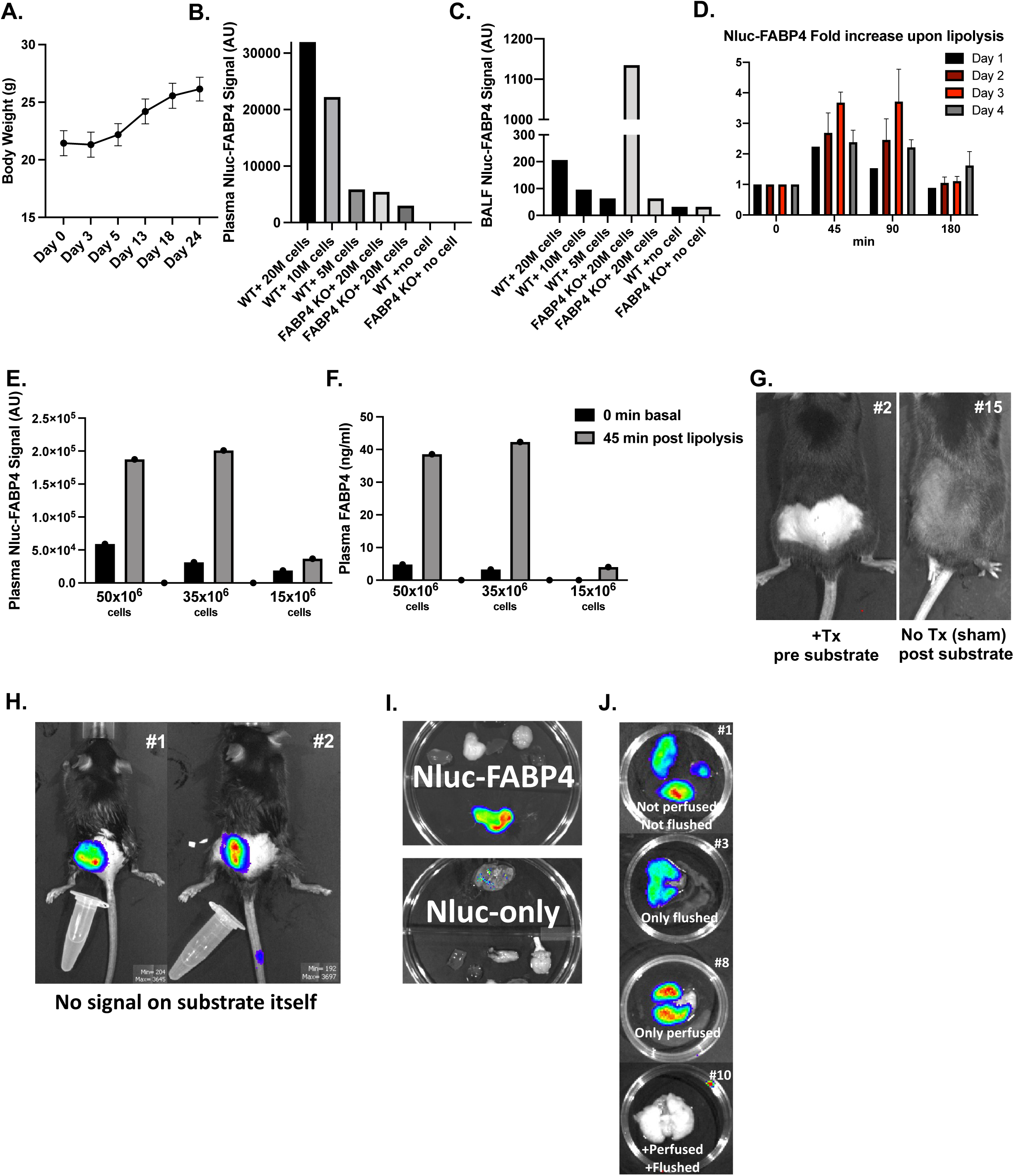
**A.** Body weight measurements of adipose tissue transplant recipient mice, starting the day of transplantation and followed up until post transplantation day 24. **B.** Plasma and **C.** BALF Nluc-FABP4 signal measured in recipient WT and FABP4 KO mice injected with Nluc-FABP4 expressing 3T3-L1 mouse adipocytes, compared to control mice injected with PBS only (no cells). **D.** Plasma time-course of Nluc-FABP4 signal fold-change following lipolysis stimulation with isoproterenol (10 mg/kg, ip) in mice injected with 50 million Nluc-FABP4 adipocytes. Implantations were made 1, 2, 3, or 4 days prior to the lipolysis experiment. Sample size: n=3 mice per group for day 2, 3, 4 experiments. **E.** Plasma Nluc-FABP4 signal upon lipolysis stimulation with isoproterenol, at baseline (0min) and 45 minutes after injection in FABP4 KO mice 3 days after implantation with the indicated numbers of Nluc-FABP4 adipocytes. **F.** Plasma FABP4 protein levels measured by ELISA using samples collected from the FABP4 KO mice shown in panel E. **G.** Negative controls in IVIS imaging. Mouse #2 was implanted with Nluc-FABP4 3T3-L1 cells but did not receive substrate injection. Mouse #15 received PBS instead of cells, both were intravenously injected with substrate 30 minutes before imaging. C57BL/6 WT mice was used for this control experiment. **H.** No luminescence signal detected from the substrate itself in microcentrifuge tubes, which were simultaneously imaged with mice (#1 and 2) that received Nluc-FABP4 3T3-L1 cells and had substrate injected 30 minutes before imaging. C57BL/6 WT mice was used for this control experiment. **I.** Nluc signal from organs and lungs collected from a mouse that received Nluc-FABP4 3T3-L1 cells (top) and a mouse that received Nluc-only 3T3-L1 cells (bottom), 30 minutes after intravenous substrate injection, followed by saline perfusion prior to tissue collection. In the top panel, the lung is positioned at the bottom of the dish, whereas in the bottom panel, the lung is positioned at the top of the dish. **J.** Detection of Nluc-FABP4 signal in lungs collected from mice implanted with Nluc-FABP4-expressing 3T3-L1 cells, 30 minutes after substrate injection via the tail vein. Panels from top to bottom show: mice #1 - Lungs collected without any saline perfusion or tracheal saline flushing, mice #3 - Lungs collected without perfusion but flushed with 5 ml saline via tracheostomy, mice #8 - Lungs perfused with 20 ml saline only, and mice #10 - Lungs perfused followed by tracheal saline flushing.

**Supplemental Figure 5.**
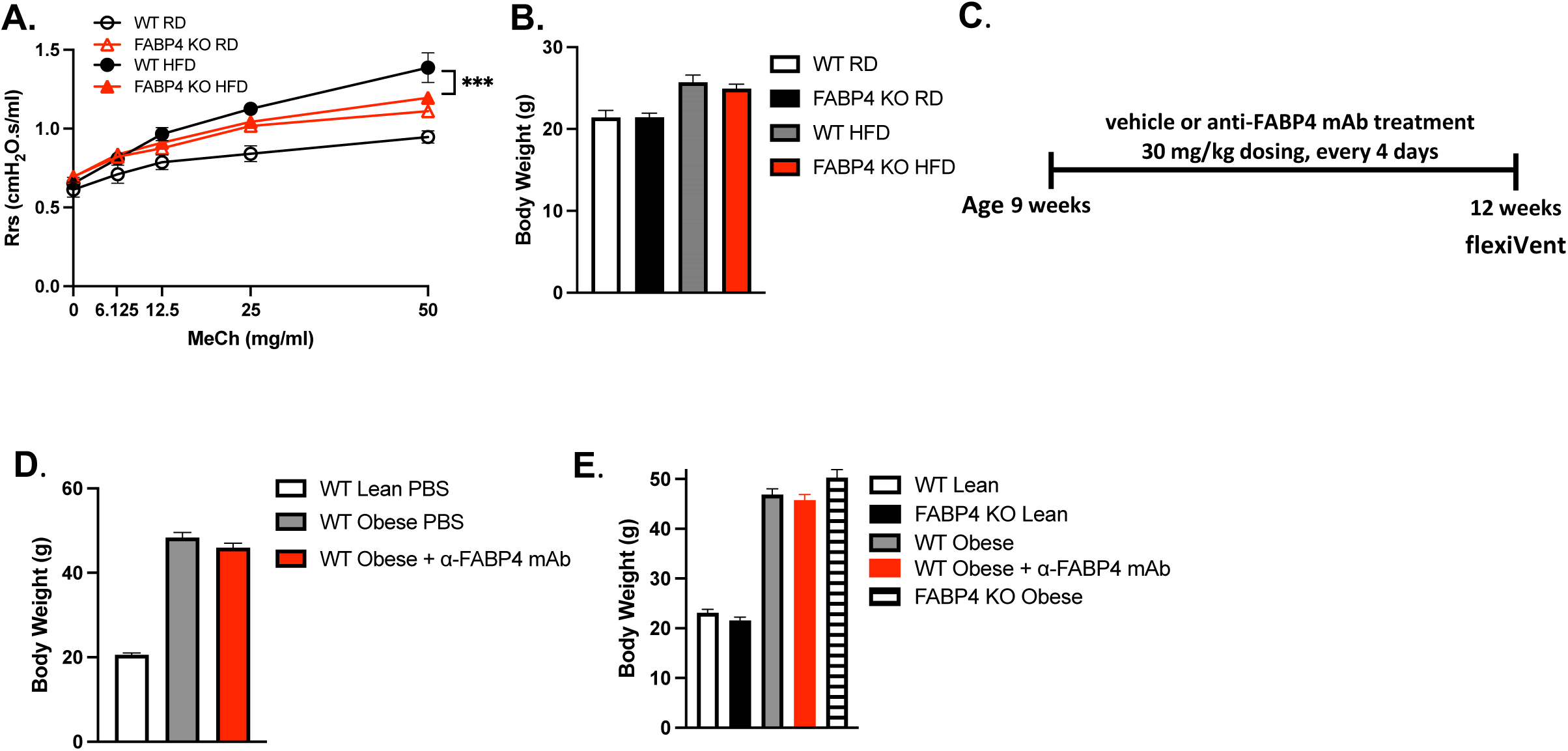
**A.** Airway hyperresponsiveness measured in WT lean (RD), WT obese (high-fat diet, HFD), FABP4 KO lean (RD), and FABP4 KO obese (HFD) mice, with sample sizes of n=3, 6, 7, and 11, respectively. **B.** Body weights of the mice shown in panel A **C.** Study design for the monoclonal antibody treatment cohort shown in figures 5D and 5G. **D.** Body weights of mice in figure 5D. **E.** Body weights of mice in figure 5G.

